# Organizational properties of a functional mammalian cis-regulome

**DOI:** 10.1101/550897

**Authors:** Virendra K. Chaudhri, Krista Dienger-Stambaugh, Zhiguo Wu, Mahesh Shrestha, Harinder Singh

## Abstract

Mammalian genomic states are distinguished by their chromatin and transcription profiles. Most genomic analyses rely on chromatin profiling to infer cis-regulomes controlling distinctive cellular states. By coupling FAIRE-seq with STARR-seq and integrating Hi-C we assemble a functional cis-regulome for activated murine B-cells. Within 55,130 accessible chromatin regions we delineate 9,989 active enhancers communicating with 7,530 promoters. The cis-regulome is dominated by long range enhancer-promoter interactions (>100kb) and complex combinatorics, implying rapid evolvability. Genes with multiple enhancers display higher rates of transcription and multi-genic enhancers manifest graded levels of H3K4me1 and H3K27ac in poised and activated states, respectively. Motif analysis of pathway-specific enhancers reveals diverse transcription factor (TF) codes controlling discrete processes. The cis-regulome strikingly enriches for combinatorial DNA binding regions of lineage determining TFs. Their genomic binding patterns reveal that onset of chromatin accessibility is associated with binding of simpler combinations whereas enhancer function requires greater complexity.

## Introduction

Metazoan cell types manifest multitudes of distinctive genomic states that are associated with their developmental history, differentiation outcome and physiological responses (Ho et al., 2014). Such genomic states can be analyzed using the framework of gene regulatory networks (GRNs) (Davidson, 2010; Thompson et al., 2015). According to this unifying framework, each stable metazoan cell state is controlled by a distinctive GRN comprised of a large set of transcription factors (TFs) and co-regulators (co-activators, co-repressors, chromatin modifiers). These in turn operate on vast but determinate sets of regulatory DNA sequences in genomes termed cis-regulomes. Changes in cell states during development or differentiation are brought about by signaling induced alterations in GRNs. Although attempts are being made to analyze the extremely complex GRNs of hundreds of distinctive mammalian cell states there remain formidable challenges. From the standpoint of analyses of TFs, rigorous analyses are proceeding with small combinations of TFs that appear to represent major determinants of cellular identities (Braun and Gautel, 2011; Martello and Smith, 2014; Singh et al., 2005; Waardenberg et al., 2014). The functions of other expressed TFs in controlling such states remain largely unexplored. Greater progress is being achieved in analyzing the diverse and intricate cis-regulomes of mammalian (murine and human) cell states. This primarily involves one or more types of chromatin profiling including histone ChIP-seq, TF ChIP-seq, FAIRE-seq, DNaseI-seq or ATAC-seq and intra-as well as inter-chromosomal contact maps (ChIA-PET, Hi-C) (Ho et al., 2014; Kieffer-Kwon et al., 2013; Rao et al., 2014). However, these approaches have a major limitation in that they infer but do not directly assess functions of regulatory elements in cis-regulomes. We address this limitation by experimentally and computationally integrating various chromatin profiling approaches including Hi-C with high throughput functional analyses using STARR-seq to assemble the complex cis-regulome of a mammalian cell state.

Recently considerable progress has been made in the development of high-throughput functional screens for enhancers (Dailey, 2015). These are based on the generation of highly complex transcriptional reporter libraries which contain thousands of putative enhancers. The reporter libraries are introduced into cells of interest by transfection and their transcriptional output is analyzed by RNA-seq. Two generalizable approaches have been used in the construction of the reporter libraries. In one case, termed massively parallel reporter assay (MPRA), sequences to be functionally tested are synthesized on microarrays and then cloned upstream of a basal promoter in a reporter plasmid along with a unique barcode. Transfection of the reporter library followed by RNA sequencing of the barcodes enables a quantitative readout of the activities of individual enhancers in the complex library (Tewhey et al., 2016). In a second approach, termed self-transcribing active regulatory region sequencing (STARR-seq), genomic regions to be functionally tested are directly cloned into a reporter plasmid downstream of the promoter and in a UTR, upstream of the polyadenylation site. Transfection of the reporter library followed by directed RNA sequencing of the enhancer sequences, which serve as their own tags, provides a quantitative readout of activities (Arnold et al., 2013). The latter approach is advantageous from the standpoint that it is more direct, not involving synthesis or tagging and it enables the functional testing of native sequences that are greater than 200bp in length. STARR-seq was developed in the context of assaying regulatory sequences in the Drosophila genome and has been deployed to a limited extent through focused screens in mammalian cells (Babbitt et al., 2015; Vockley et al., 2016). We couple formaldehyde-assisted isolation of regulatory elements (FAIRE-seq) with STARR-seq thereby enabling direct functional assessment of a diverse and complex set of accessible chromatin regions for enhancer activity. Importantly, by computationally integrating this data with chromatin interactions mapped by Hi-C we are able to assemble a prototype functionally-filtered cis regulome for a primary mammalian differentiated cell state.

B-lymphocytes of the mammalian immune system have served as a leading model for the analysis of both cell type-specific and activation induced GRNs (Cobaleda et al., 2007; Grosschedl, 2013; Kieffer-Kwon et al., 2013; Mansson et al., 2012; Xu et al., 2015). Furthermore, a diverse array of chromatin profiling approaches, have been used to delineate the cis-regulomes of resting and bacterial lipopolysaccharide (LPS) activated murine splenic B-cells and Epstein-Barr virus transformed human B-cell lines, the latter as a key part of the ENCODE project (Thurman et al., 2012). However, as noted above none of these approaches have involved comprehensive functional screens of the large numbers of presumptive transcriptional enhancers identified by chromatin profiling. We build a cis-regulome for LPS activated murine splenic B-cells by genomic integration of structural (FAIRE-seq), functional (STARR-seq) and connectivity (Hi-C) datasets. These are superimposed with H3K27Ac, H3K4Me3 and TF ChIP-seq profiles to reveal 9,989 enhancers, including 29 super-enhancers, which are connected to promoters of 7,530 actively transcribed genes. The cis-regulome is dominated by long range enhancer-promoter interactions (>100kb) and their complex combinations. Genes with multiple enhancers display higher rates of transcription and multi-genic enhancers manifest higher levels of H3K4me1 and H3K27ac in their poised and activated states, respectively. The functional cis-regulome in contrast with accessible chromatin regions is enriched for combinatorial sets of DNA bound TFs including PU.1, E2A, Ebf1, Pax5 and Blimp1, known to control the activated B-cell state. Motif analysis of enhancers linked to biologically coherent gene modules implicates additional TFs and diverse combinatorial codes in the control of discrete molecular pathways and processes.

## Results

### Functional filtering of enhancers by coupling FAIRE-seq with STARR-seq

To build a cis-regulome of a mammalian cellular state one needs to acquire four types of genomic information (i) the set of actively transcribed genes, (ii) the set of accessible chromatin regions that includes those marked with activating histone modifications and representing poised or active enhancers, (iii) the set of functional enhancers, and (iv) the contacts of functional enhancers with active promoters. This goal can be achieved through the experimental and computational integration of RNA-seq, FAIRE-seq, STARR-seq and Hi-C datasets (Figure 1A). To profile and capture accessible chromatin regions for functional filtering, we reasoned that FAIRE-seq would be advantageous. This chromatin profiling approach enriches for short DNA fragments (100-400 bp, lengths of typical enhancers) that are TF-bound but nucleosome-depleted (Gaulton et al., 2010). Furthermore, these fragments can be cloned into a STARR-seq vector enabling their high-throughput functional screening (Figure 1A).

**Figure 1.**
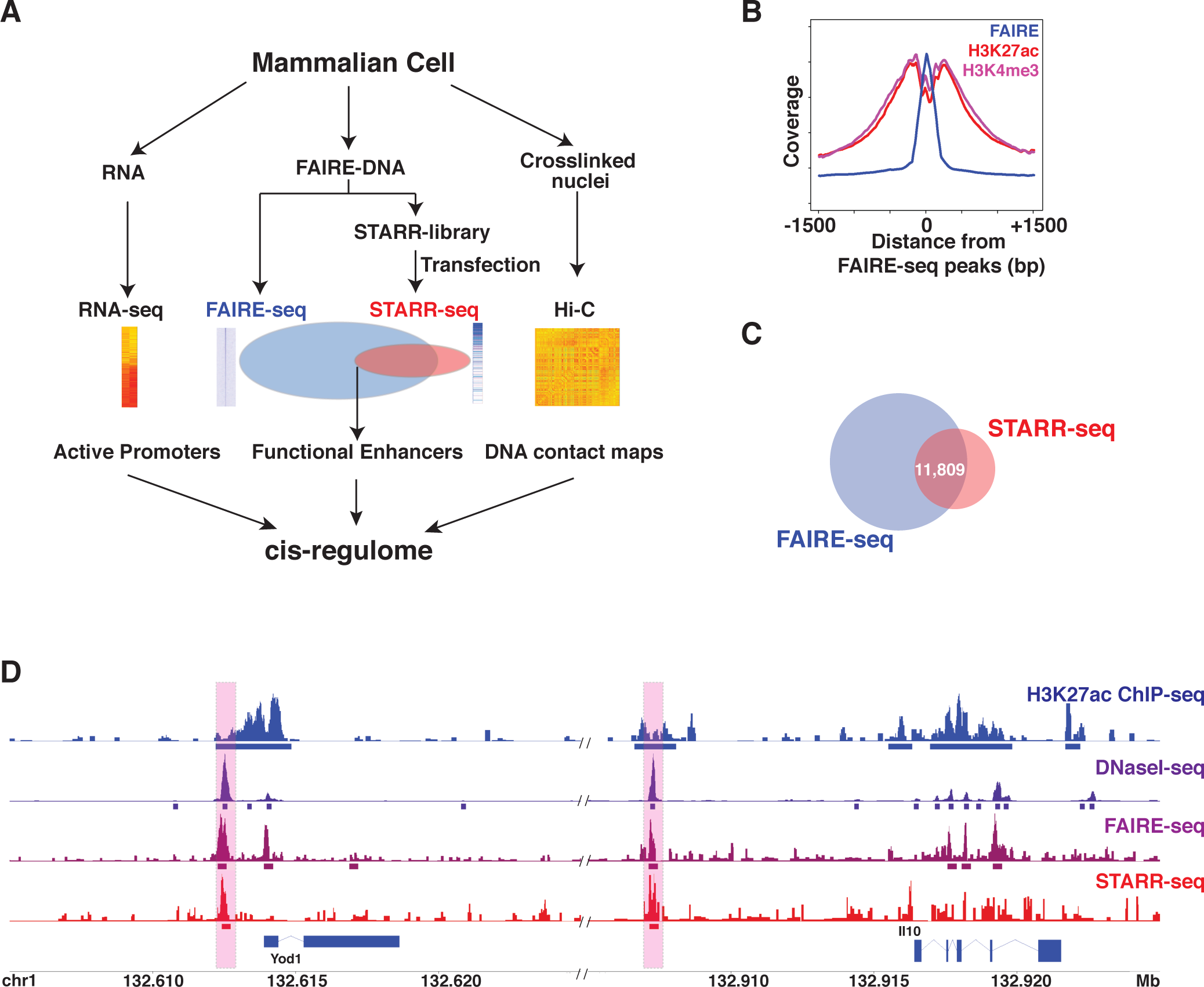
Assembling a functional cis-regulome for a mammalian cell state **A.** Schematic illustrates distinct genome-wide structural (FAIRE-seq), functional (STARR-seq) and connectivity (Hi-C) assays that are deployed and computationally integrated with RNA-seq to assemble a cis-regulome for a mammalian cell state. In this experimental design, FAIRE-seq DNA fragments which reflect open chromatin regions are cloned into a STARR-seq reporter vector and the resulting plasmid library is transfected into relevant mammalian cells and screened for enhancers. **B-D.** Purified splenic murine B-cells, activated with LPS for 72 hours, were used for all genomic assays**. B.** Formaldehyde cross-linked chromatin from activated B-cells was processed for FAIRE-seq, H3K27ac and H3K4me3 ChIP-seq. Averaged tag densities using a 10 bp bin size are plotted after normalization across all FAIRE-seq peak regions along with an overlay of their H3K27ac and H3K4me3 profiles. **C.** Overlap of STARR-seq peaks with FAIRE-seq peaks. STARR-seq plasmid library, comprising of size-selected FAIRE DNA fragments, was transfected into LPS activated B-cells (48 or 60h post-activation) and RNA was isolated for sequencing of reporter transcripts at 72h. Peaks were called using HOMER pipeline. **D.** A genome plot of the IL-10 locus displaying the H3K27ac ChIP-seq, DNaseI-seq, FAIRE-seq and STARR-seq tracks. HOMER assigned peaks are indicated with corresponding color-coded bars below the tracks. Two enhancers identified by STARR-seq that reside upstream of the IL-10 promoter in accessible chromatin regions marked by H3K27Ac are highlighted. A genome scale bar with the IL-10 and Yod1 loci is shown below.

For all genomic analyses, we used purified murine splenic B-cells that were activated with LPS for 72h. This experimental system has been extensively used in numerous genomic studies to analyze the activated B-cell state (Kieffer-Kwon et al., 2013; Xu et al., 2015). Upon stimulation with LPS, naïve, resting B-cells exit G_0_, undergo metabolic re-programming and enter S phase at 24h (Turner et al., 2008). The cells then undertake 2-4 cell divisions with a fraction of the progeny (15-20%) differentiating into immunoglobulin (Ig) secreting plasmablasts by 72h (Xu et al., 2015). LPS-activated B-cells were profiled using RNA-seq and FAIRE-seq along with ChIP-seq for H3K27Ac and H3K4me3 (see <u>UCSC browser link)</u> (Supplementary Figures S1A-D). The latter two genomic analyses enabled us to determine if a large number of the FAIRE-seq peaks were positioned between adjacent nucleosomes that are marked by the histone modifications, H3K27Ac and/or H3K4me3. Indeed FAIRE-seq peaks with a width of 300bp were seen to reside between nucleosome peaks bearing the activating histone modifications (Figure 1B). Thus, the FAIRE-seq DNA fragments from LPS activated B-cells, representing 55,133 peaks, were deemed to be a comprehensive and suitable source for a high-throughput functional screen of enhancers using STARR-seq. Importantly, approximately 45,000 of our FAIRE-seq peaks were additionally validated by aligning them with independently generated DNaseI-seq profiles of activated B-cells (Kieffer-Kwon et al., 2013) (see <u>UCSC browser link</u>). The FAIRE-seq DNA fragments were cloned into the mammalian STARR-seq vector between the GFP gene and the SV40 poly(A) signal sequence (Arnold et al., 2013). Sequencing of the plasmid library revealed >81% coverage of the input FAIRE-DNA, thereby generating a representation of 44,560 FAIRE-seq peak regions analyzed above. The STARR-seq plasmid library was transfected into activated B-cells at either 48h or 60h post LPS activation and RNA was isolated for deep sequencing of plasmid transcripts at 72h (see Methods). Data from biological replicates revealed excellent reproducibility at both transfection times (Supplementary Figures S1E, F). The 60h transfection assay time was used to delineate enhancers that are functional at late times after B-cell activation as the cells are undergoing a bifurcation with a subset differentiating into plasmablasts and others into pre-germinal center (GC) B-cells (Xu et al., 2015). Given that activated B-cells undergo considerable cell death after 72h of LPS stimulation we harvested the cells for RNA isolation at that timepoint. Peaks were called in the STARR-seq datasets using HOMER after merging the replicate 12h or 24h datasets. The peak regions from the two datasets were then combined to constitute the set of functional regions for further analysis. Intersection of the STARR-seq peaks with the FAIRE-seq peaks revealed 11,809 regions that overlapped (Figure 1C). It should be noted that FAIRE-seq and STARR-seq peaks (300 bp) were called with HOMER using the same parameters. As anticipated, the different transfection assay timepoints identified both shared (7,089) as well as unique STARR-seq peaks (4,720). The biological significance of the shared versus kinetically distinguishable enhancers is elaborated below. We note that 5,570 STARR-seq peaks in our transfection assay did not overlap with FAIRE-seq peaks (Figure 1C). The detection of functional enhancers in poorly accessible chromatin regions has been noted in the initial STARR-seq study (Arnold et al., 2013). These regions could represent enhancers that are dependent on prior chromatin remodeling for their function. However, we excluded them in the subsequently assembly of the cis-regulome as they lacked a key chromatin feature. Nevertheless, coupling of FAIRE-seq with STARR-seq enabled efficient genome-wide functional filtering of accessible regions in mammalian chromatin for enhancer activity. An alignment of the H3K27Ac, DNaseI-seq, FAIRE-seq and STARR-seq genome tracks for the IL-10 locus is shown in Figure 1D (see also <u>UCSC browser link</u>). Importantly, in activated B-cells, approximately one-fifth of FAIRE-seq peaks, that are also DNaseI accessible, functioned as enhancers in the STARR-seq assay.

### Validation of STARR-seq enhancers

As anticipated the largest number of STARR-seq enhancers were positioned within intergenic regions with the remaining distributed in promoter proximal or intragenic locations (Figure 2A). To further substantiate the B-cell enhancers identified by STARR-seq we undertook both genome-wide analyses as well as an independent functional sampling approach. Active enhancers have been shown to bind the transcriptional co-activator p300 (Visel et al., 2009). Based on alignment with an available p300 ChIP-seq dataset in activated B-cells (Kieffer-Kwon et al., 2013), nearly half of the STARR-seq enhancers were bound by p300 (Figure 2B). Importantly, p300 binding was more strongly enriched in STARR-seq peaks in relation to FAIRE-seq peaks. A similar fraction of the STARR-seq peaks were also enriched for the binding of CTCF in activated B-cells (Figure 2B) (Nakahashi et al., 2013) a transcription factor which has been shown to promote long-range interactions between enhancers and promoters (Ren et al., 2017). Transcriptional enhancers have also been shown to generate enhancer-derived transcripts termed eRNAs (Palozola et al., 2017). We used pulse labeling of nascent transcripts with the uridine analog EU and Click-iT chemistry (Palozola et al., 2017) to purify and sequence nuclear RNA from activated B-cells (Figure 2C). We confined our analysis to intergenic FAIRE-seq accessible regions so as to focus on non-coding transcripts. Mapping of such nascent transcripts to FAIRE-seq accessible regions that did or did not overlap with a STARR-seq peak revealed STARR-seq positive regions to be strongly enriched for eRNAs (p=1.0e-118) (see also <u>UCSC browser</u> <u>link</u>).

**Figure 2.**
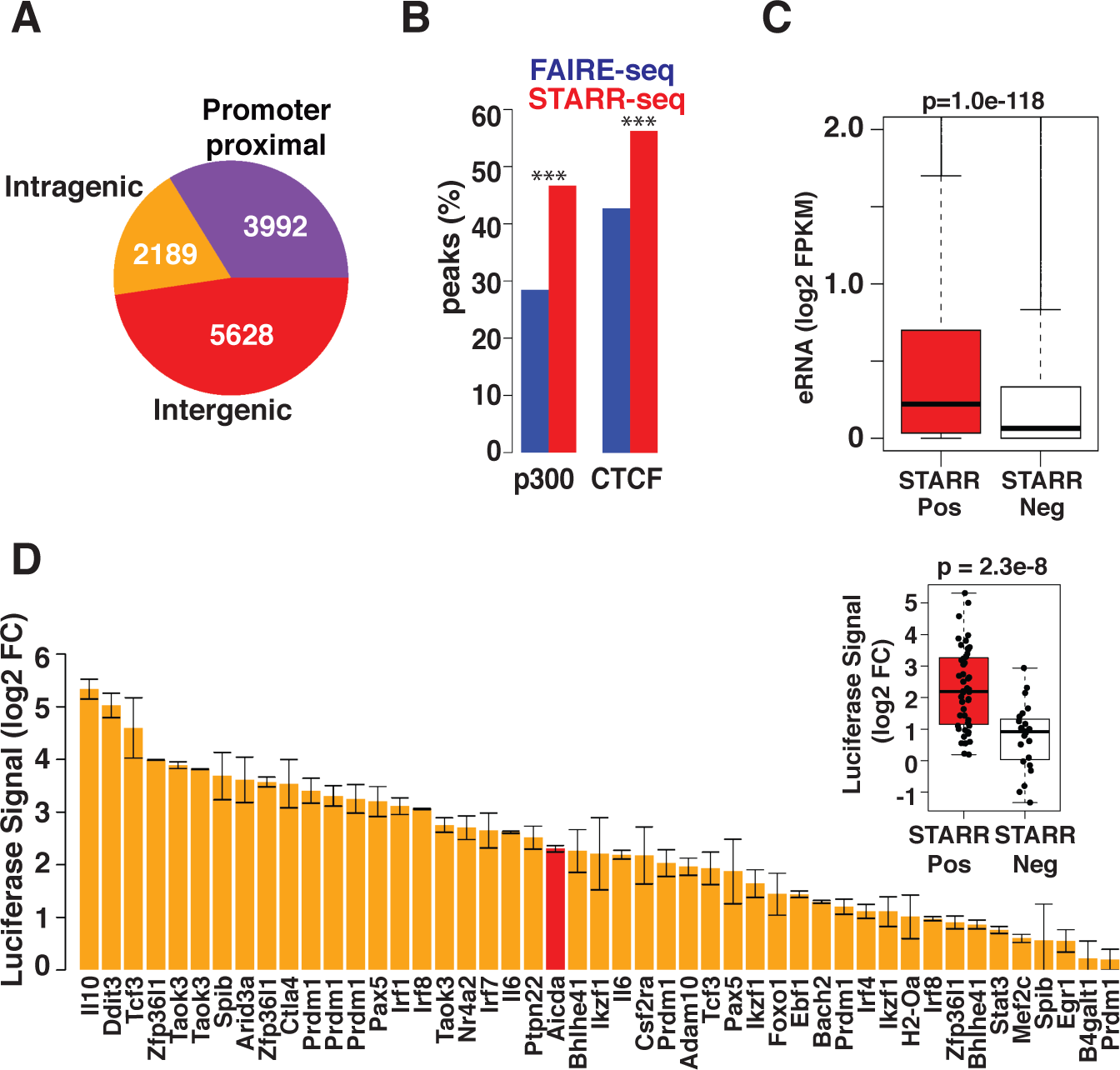
Validation of STARR-seq identified enhancers **A.** Genomic distribution of STARR-seq identified enhancers (n=11,809). Promoter proximal enhancers overlap with regions around transcriptional start sites (−1kb to +100bp). **B.** Histogram displaying the percentages of FAIRE-seq and STARR-seq peaks that overlap with ChIP-seq peaks for p300 and CTCF in activated murine B-cells (Kieffer-Kwon et al., 2013; Nakahashi et al., 2013) STARR-seq enhancers more strongly enrich for binding of P300 and CTCF as compared to FAIRE-seq peaks (*** denote p-values < 1.0e-20 using Fisher’s exact test). **C.** Analysis of enhancer associated RNAs (eRNA) spanning STARR-seq positive (STARR Pos) versus STARR-seq negative (STARR Neg) peaks. eRNAs were detected by pulse labeling of nascent transcripts in activated B-cells with the uridine analog EU and quantified (FPKM) by extending a 300 bp window on each side of all intergenic FAIRE-seq peaks that were either STARR-seq peaks (STARR Pos) or not (STARR Neg). The control set of sequences (STARR Neg) were positioned at least 1kb away from a STARR Pos region and did not overlap with H3K27Ac domains containing one or more enhancers. p-value is Kolmogorov-Smirnov test (KS test). **D.** Validation of select enhancers in CH12 B-cell line. STARR-seq enhancers (n=47) identified in a subset of biologically important B-cell genes were cloned into a luciferase reporter vector and assayed by transfection in CH12 cells. The bar plot displays distribution of luciferase reporter signals after normalization with empty vector for the indicated gene enhancers. The Aicda enhancer (red bar) was included as a positive control. The box plot shows the distribution of enhancer activity for the 47 STARR-seq positive regions in panel **E** compared with 23 STARR-seq negative regions. The latter also corresponded with FAIRE-seq peaks. p-value is for t-test.

To complement the above analysis, we sampled a set of STARR-seq enhancers (n=46) by individually assessing their functionality using a luciferase reporter system (Figure 2D). These enhancers were selected based on their positioning near biologically important B-cell genes. To generate a control set of DNA sequences we functionally sampled FAIRE-seq positive regions which did not generate a STARR-seq peak. A known enhancer from the *Aicda* gene was used a positive control. The reporter assays were performed in the murine CH12 B-cell line, which has been extensively used as a model for the activated B-cell state and plasmablast differentiation (Kieffer-Kwon et al., 2013). The B-cell line enabled much higher sensitivity of detection of luciferase expression with transfected reporter constructs. Of the 46 enhancers tested in CH12 cells, 38 (~83%) exhibited 2-fold or greater enhancement of reporter gene expression (Figure 2D). In contrast, the control set of STARR-seq negative DNA sequences (n=23) displayed significantly lower activity than their STARR-seq positive counterparts (Figure 2D, inset). Thus, the STARR-seq identified enhancers were corroborated using both genome-wide molecular features and independent functional sampling.

### Using Hi-C to complete assembly of B-cell cis-regulome

FAIRE-seq coupled with STARR-seq delineated 11,809 enhancers. Given that enhancers can activate transcription from cognate promoters at highly variable genomic distances we performed high resolution chromosome conformation capture (Hi-C) (Rao et al., 2014) to connect the functional enhancers with active promoters in B-cells. Importantly, this approach utilized unbiased genome-wide contact maps to compile all intra-chromosomal interactions (Supplementary Figure S2). A 100kb resolution contact map for normalized interactions on mouse chromosome 10 is shown (Figure 3A). Higher resolution contact maps were then used to identify enhancer-promoter pairs that were contained within interacting bins of varying sizes (10, 5, 2 and 1kb) so as to capture both long- and short-range interactions. Active promoters were delineated using the RNA-seq dataset (Supplementary data S1) and represented those of expressed genes whose transcript levels were above a threshold of 4 FPKM (8,199 genes). Enhancer-promoter interactions over 10 Mb were not considered. To connect enhancers with promoters we defined promoter regions to be within −1000 to +100bp of the transcriptional start site and interacting pairs were identified based on overlap between promoter and enhancer bins. Only direct enhancer-promoter contacts were included in the analysis. We thus connected 9,989 enhancers (84% of STARR-seq enhancers) to promoters of 7,530 genes (92% of expressed genes) (Supplementary data S2, B-cell cis-regulome resource) (see also <u>UCSC browser link</u>). As an example, high resolution contact map (5kb) for the *Prdm1* locus, which encodes the TF Blimp1, a regulator of plasma cell differentiation, is shown (Figure 3B). The locus was observed to have 8 long-range enhancers interacting with its promoter, ranging in distance from 147 to 585Kb (Figure 3B). To contextualize this finding, we analyzed the distribution of genomic distances for all enhancer-promoter contacts (Figure 3C). The median distance was 495kb, considerably larger than that of all Hi-C interactions (172kb). Thus, the genomic distances exemplified by the *Prdm1* enhancers lie at or below the median. Based on our tri-partite workflow, that involves integration of chromatin profiling, a functional screen and Hi-C, we refer to the compilation of the 9,989 enhancers as the assembled B-cell cis-regulome. This framework is readily generalizable to other cellular contexts.

**Figure 3.**
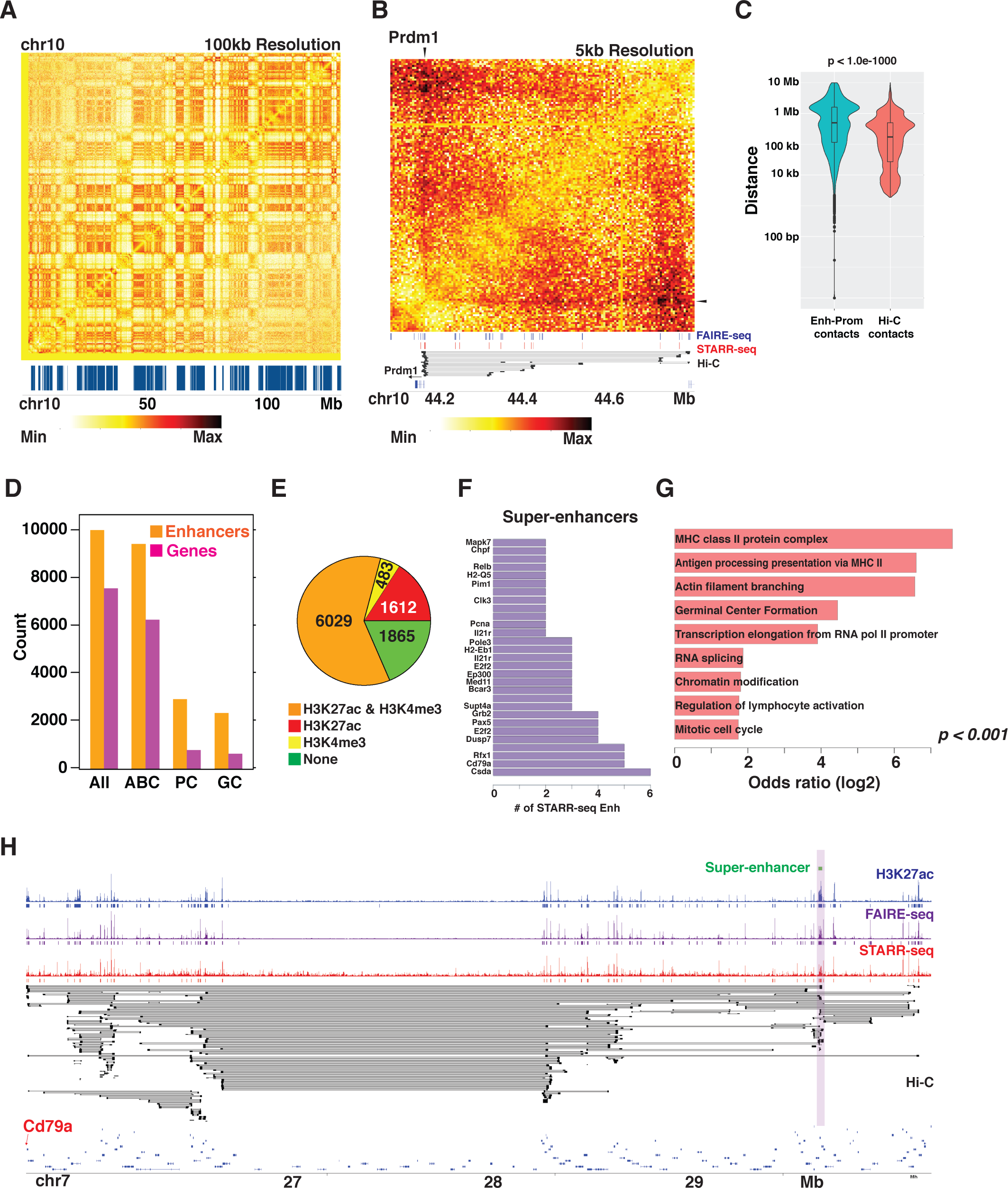
Generating a connectivity map of B-cell enhancers and active promoters using Hi-C Hi-C was performed with nuclei isolated from formaldehdye cross-linked, activated B-cells, using the restriction endonuclease MboI, thereby generating a high-resolution genomic contacts matrix. **A.** Hi-C contact matrix for murine chromosome 10. Heatmap displays normalized contact read counts at a resolution of 100 kb bin size. Hi-C compartments are displayed with blue bars. **B.** A higher resolution contact matrix (5 kb bin size) of the *Prdm1* locus on chromosome 10. All enhancer-promoter connectivity links for the *Prdm1* locus in the Hi-C dataset along with their bin sizes (10 or 5 kb) are shown below the contact matrix (black bars). FAIRE-seq and STARR-seq peaks are displayed above the connectivity links. **C.** Violin plots of the distances and relative frequencies of enhancer-promoter contacts in the B-cell cis-regulome. This distribution is compared with that of all intra-chromosomal interactions in the Hi-C dataset. P-value is based on KS test. **D.** Numbers and types of genes whose promoters are connected with functional enhancers by Hi-C in B-cell cis-regulome. ABC refers to genes expressed in activated B-cells. PC and GC refer to genes that are preferentially expressed in plasma cells and germinal center B-cells, respectively. **E.** Pie chart of cis-regulome enhancers embedded in H3K27ac and H3K4me3 peaks. Coordinates of H3K27Ac and H3K4me3 ChIP-seq peaks were overlapped with those of STARR-seq peaks for cis-regulome enhancers to generate the frequency distribution. **F.** Enumeration of super-enhancers in cis-regulome on the basis of extended H3K27Ac ChIP-seq peak signals and the presence of 2 or more enhancers (STARR-seq peaks). Each bar represents a super-enhancer with its number of STARR-seq identified enhancers. Select biologically important genes connected to these super-enhancers are indicated. **G.** GO category enrichment analysis of genes connected with super enhancers in B-cell cis-regulome. Odds ratios for top scoring GO terms are plotted. **H.** Super-enhancer connected to Cd79a locus. Genome plot displays the H3K27ac ChIP-seq, FAIRE-seq and STARR-seq tracks. HOMER assigned peaks are indicated with corresponding color-coded bars below the tracks. All enhancer-promoter connectivity links from the Hi-C dataset along with their bin sizes (10, 5, 2 or 1kb) are displayed (black bars). The super enhancer is highlighted. A genome scale bar with the Cd79a locus on the left end is shown below.

### Characteristics of enhancers in B-cell cis-regulome

The enhancers in the assembled B-cell cis-regulome nearly completely overlapped with DNaseI accessible regions in murine B-cells and splenocytes (Supplementary Figure S3). Their overlap diminished when comparing with DNaseI accessible regions in other cellular contexts and tissues. This is consistent with B-cell-specific enhancers within the cis-regulome that manifest tissue-specific DNaseI accessibility. Analysis of the genes in the cis-regulome on the basis of their expression pattern revealed the majority to be reflective of the activated B-cell state (ABC) with a smaller number that were preferentially expressed in plasma cells (PC) or germinal center (GC) B-cells (Figure 3D). We note that the ABC genes are primarily involved in metabolic programming, cell cycle entry and mitosis.

Overlay of enhancers in the activated B-cell cis-regulome with H3K27Ac and H3K4me3 ChIP-seq profiles showed that the majority of the enhancers were covered by one or both of these activating marks (Figure 3E). Extended domains of H3K27 acetylation have been characterized as regions of high-level transcriptional activity and to span super-enhancers (Dowen et al., 2014; Hnisz et al., 2013; Whyte et al., 2013). Given that our analysis included a genome-wide functional screen of enhancers, we could uniquely assess if super-enhancers delineated on the basis of extended H3K27ac domains contained 2 or more active enhancers, that may represent their functional cores. We first identified H3K27ac super-enhancers from ChIP-seq data using conventional criteria (Whyte et al., 2013) and then screened for those that contained multiple functional enhancers. Thus, we identified 29 out of 47 extended H3K27ac demarcated super-enhancers in our datasets that contained 2 or more active enhancers (Figure 3F). Genes connected with these enhancers were strongly enriched for functions in antigen processing and presentation including actin branching that are crucial for B-T cell interactions (Figure 3G, Supplementary data S3). A super enhancer communicating with the Cd79a gene promoter, across a 3 MB span, is shown in Figure 3H. This B-cell identity gene encodes a signal transducer that is a key component of the B-cell antigen receptor and is required for B-cell development and activation. The integrated genomic analysis provides an additional and an important functional criterion for the delineation of super-enhancers. Thus, the activated B-cell cis-regulome appears to be dominated by very long-range enhancer-promoter interactions (> 100kb) including those involving functional super-enhancers regulating genes involved in lymphocyte activation and antigen presentation.

### Organizational features of B-cell cis-regulome

The assembled cis-regulome represents a comprehensive compilation of functional regulatory elements within an activated B-cell. It has been proposed that the majority of enhancers of a signal regulated cell state are poised by TF binding and chromatin remodeling reflected by H3K4me1 before their signaling-dependent activation (Heintzman et al., 2007; Heinz et al., 2010). To test this proposition, we analyzed the degree of H3K4me1 and H3K27ac in enhancers of activated B-cells before and after LPS stimulation. As predicted for their poised states these enhancer regions manifested appreciable levels of H3K4me1 in resting B-cells which diminished upon cellular activation (Figure 4A). Importantly, the reduction in the H3K4me1 levels was not observed in control FAIRE-seq regions. As expected, activation of B-cells was accompanied by increased H3K27ac levels in the cis-regulome. Thus, the activated B-cell cis-regulome enriches for poised enhancers that appear to be functionally activated in a signaling dependent manner.

**Figure 4.**
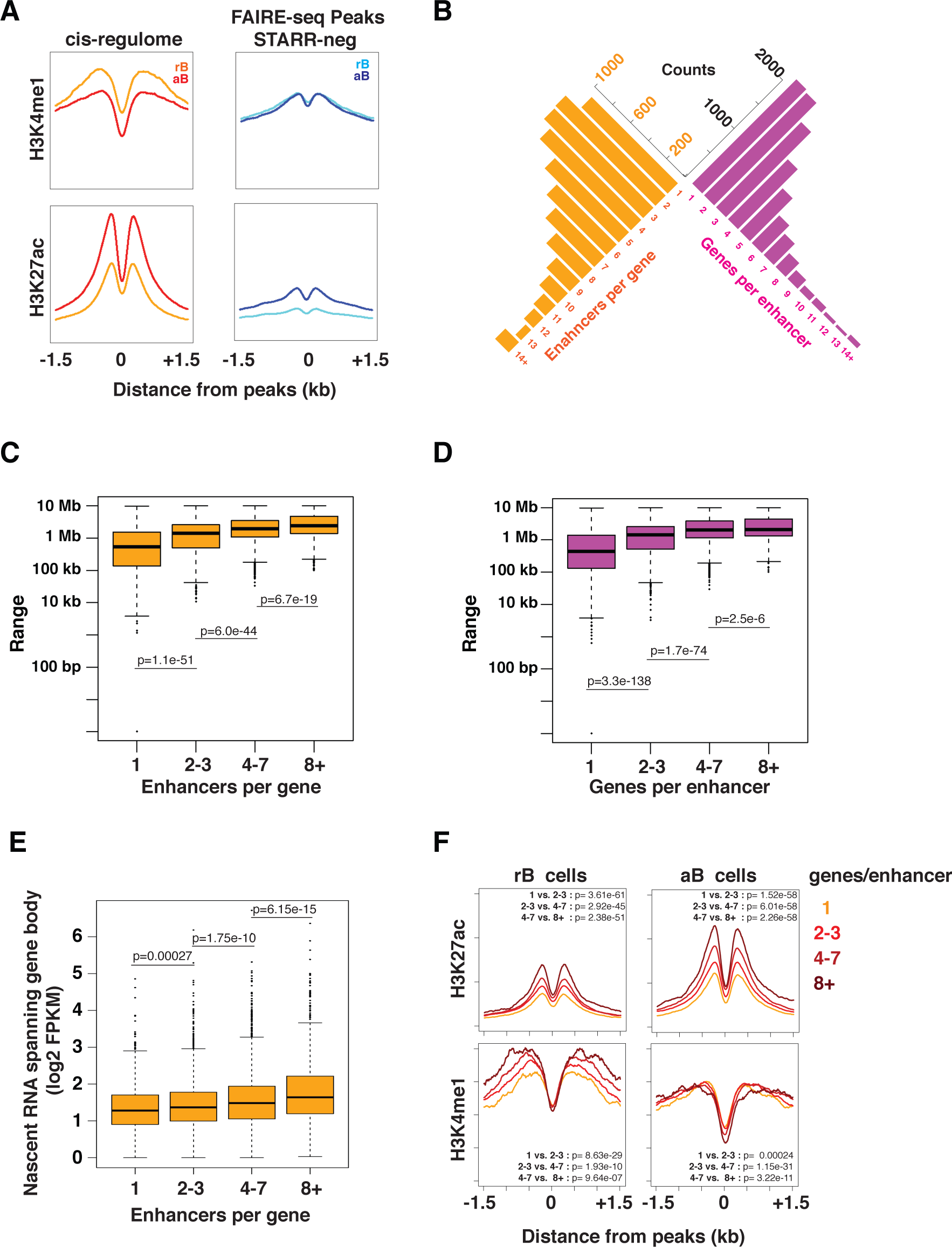
Analyzing organizational features of cis-regulome **A.** Comparison of averaged H3K4me1 and H3K27Ac tag densities for cis-regulome in resting B (rB) versus activated B (aB) cells. This encompassed 9989 regions that were FAIRE-Seq and STARR-seq peaks and connected to promoters of transcribed genes in activated B-cells. The control set of sequences were FAIRE-seq peaks (27297) that did not generate peaks in the STARR-seq assay (STARR Neg). Averaged tag densities using a 10 bp bin size are plotted after normalization. Decrease in H3K4me1 levels in cis-regulome upon B-cell activation manifested a p-value of 7.44 x 10^-48^ based on t-test. This was accompanied with an increase in H3K27ac levels with a p-value of 1.39 x 10^-32^. When comparing cis-regulome with control sequences the change in H3K4me1 (p-value 1.13 x 10^-25^) was significantly different than for change in H3K27ac (p value = 0.81), although the overall magnitude of H3K27Ac in cis-regulome was much higher. The paired t-tests were performed using coordinates flanking STARR-seq or FAIRE-seq peaks (−300 to −150 and +150 to +300). **B.** Frequency distributions enumerating connectivity of enhancers and genes in B-cell cis-regulome. Counts of enhancers associated per gene (orange bars) and promoters of genes associated with individual enhancers (magenta bars) are displayed. **C.** Boxplot for range of action for enhancers binned for the number of genes they act upon. **D.** Boxplot showing the distance to farthest enhancer acting on a gene where genes are binned based on number of enhancers acting upon them. **E.** Quantitative analysis of nascent transcripts, enumerated by EU labeling, across gene bodies in relation to the numbers of enhancers acting on promoters of genes. **F.** H3K27ac and H3K4me1 ChIP-seq coverage plot for the enhancers binned as in D, showing histone modification as a function of number of genes that enhancers are connected to. Indicated p-values for panels **C-F** are based on t-test. In panel F, the paired t-tests were performed using coordinates flanking STARR-seq peaks (−450 to −150 and +150 to +450).

Next, we examined the frequency distributions of connectivity between genes and enhancers by examining the numbers of enhancers per gene (orange) as well as genes per enhancer (purple) (Figure 4B). The vast majority of genes were (6572) were observed to be connected with 2 or more enhancers. In this regard, the *Prdm1* locus is representative of a large number of genes in the activated B-cell cis-regulome (1312, ~17% of total) that are communicating with 8 or more enhancers. Conversely, most enhancers (7840) were interacting with the promoters of 2 or more genes. Surprisingly, ~40% of the cis-regulome involved individual enhancers interacting with promoters of 4 or more genes. Given these two types of frequency distributions, enhancers per gene and genes per enhancer, we analyzed the maximal range (distance) of action of enhancers as a function of the numbers of enhancer-gene pairs (Figure 4C, D). This analysis revealed that an increase in the number of enhancer-gene interactions for either distribution was associated with an increased range of action for enhancers. Thus, multiple enhancers acting on a promoter result in extension of the range of action of the distal most enhancer. Similarly, the range of action of an enhancer increases with the number of promoters that it is interacting with.

Given the finding of multiple enhancers communicating with genes in the cis-regulome we examined the relationship between the number of active enhancers that were interacting with the promoter of a given gene and the magnitude of nascent transcripts across the gene body. This was experimentally determined by EU labeling (Palozola et al., 2017) (Figure 4E). The analysis showed that an increase in active enhancers per gene was strongly correlated with higher levels of transcription, consistent with a cumulative mechanism of control. Next, we analyzed the relationship between the number of genes (promoters) an enhancer was communicating with and its H3K27ac levels. Intriguingly, H3K27ac levels of enhancers in both resting as well as activated B-cells showed a strong correspondence with the number of genes that the enhancers were communicating with (Figure 4F). There was a proportional increase in enhancer H3K27ac levels as the numbers of interacting promoters increased. Surprisingly, there was a similar relationship between enhancer H3K4me1 levels and the numbers of cognate promoters in resting B-cells which however was greatly diminished upon B-cell activation. Thus, the activated B-cell cis-regulome appears to be dominated by long range and complex enhancer-promoter interactions. This organization encompasses multiple enhancers acting on a given gene from large distances that appear to cumulatively control the magnitude of transcription. It also reveals enhancers interacting with multiple promoters. This results in a gradation of poised and activated states for such enhancers that are associated with the number of interacting promoters.

### TF motif analysis of enhancers – control of functionally coherent gene modules

To gain insight into the transcription factors that are acting on the B-cell cis-regulome we tested for the enrichment of TF binding site motifs using HOMER. The TF motif database utilized by HOMER was supplemented with TF motifs for all expressed B-cell TFs that were annotated in CIS-BP (Weirauch et al., 2014) (Supplementary data S4). Motif enrichment in the test set of cis-regulome sequences (accessible chromatin, STARR-seq peaks that contact active promoters) was carried out by referencing to a scrambled set of sequences (background set) that maintained the same base pair composition and di-as well as tri-nucleotide frequencies. This analysis revealed 45 unique TF motifs that were enriched by 2-fold or greater (log p-value < −20, Supplementary data S5). Importantly, the motifs for the B-lineage determining TFs PU.1(EICE), E2A (Tcf3), Ebf1 and Blimp1 (Prdm1) were identified in this set (Figure 5A). Furthermore, the motifs for canonical signaling-induced TFs such as NF-κB, NFAT, AP-1 and STATs, known to function in B-cell activation were also seen to be strongly enriched in the B-cell cis-regulome. Next, we sought to delineate TF motifs that correlated with particular enhancer features. For this analysis we used STARR-seq selected tags associated with the cis-regulome, which enriched for functional sequences within the enhancer coordinates. Analysis of enhancer tags captured at the two different time points for transfection of the STARR-seq library revealed selective enrichment of certain TF motifs (Supplementary Figure S4, Supplementary data S6). In particular motifs for TFs such as Bach2, Ets1 and Pax5 that antagonize plasma cell differentiation were enriched in the late time point. We next tested whether certain TF motifs were preferentially enriched in the context of proximal versus distal enhancers or in super-enhancers. Comparison of proximal (−1kb to +100bp) versus distal enhancers revealed differential enrichment of the B-lineage-determining TF motifs with the Pax5 motif in the former and the E2A (TCF3), Ebf1, PU.1 (EICE) and Prdm1 motifs in the latter (Figure 5A, Supplementary data S7). Intriguingly, the B-cell super-enhancers when compared to rest of the cis-regulome were enriched for the Tcf3 and Pou2f1 motifs, the latter is recognized by Oct-1 and Oct-2 along with OCA-B (Figure 5B). Oct-2 and OCA-B are selectively expressed in activated B-cells and play important roles in controlling B-cell responses (Kim et al., 1996). Furthermore, B-cell super-enhancers revealed strong enrichment for motifs recognized by the signaling induced TFs Srf, NF-κB, NFAT and Stat6. Thus, the enhancers of the B-cell cis-regulome contain motifs for the major B-cell determining TFs as well as the more broadly acting signaling induced TFs. The latter are known to be activated by signaling through the B-cell receptor (BCR), Toll-like receptors (TLR) as well as co-stimulatory (CD40) and cytokine receptors, expressed on the surface of B-cells and required for their proliferation and differentiation in distinct immune responses (Kurosaki et al., 2010).

**Figure 5.**
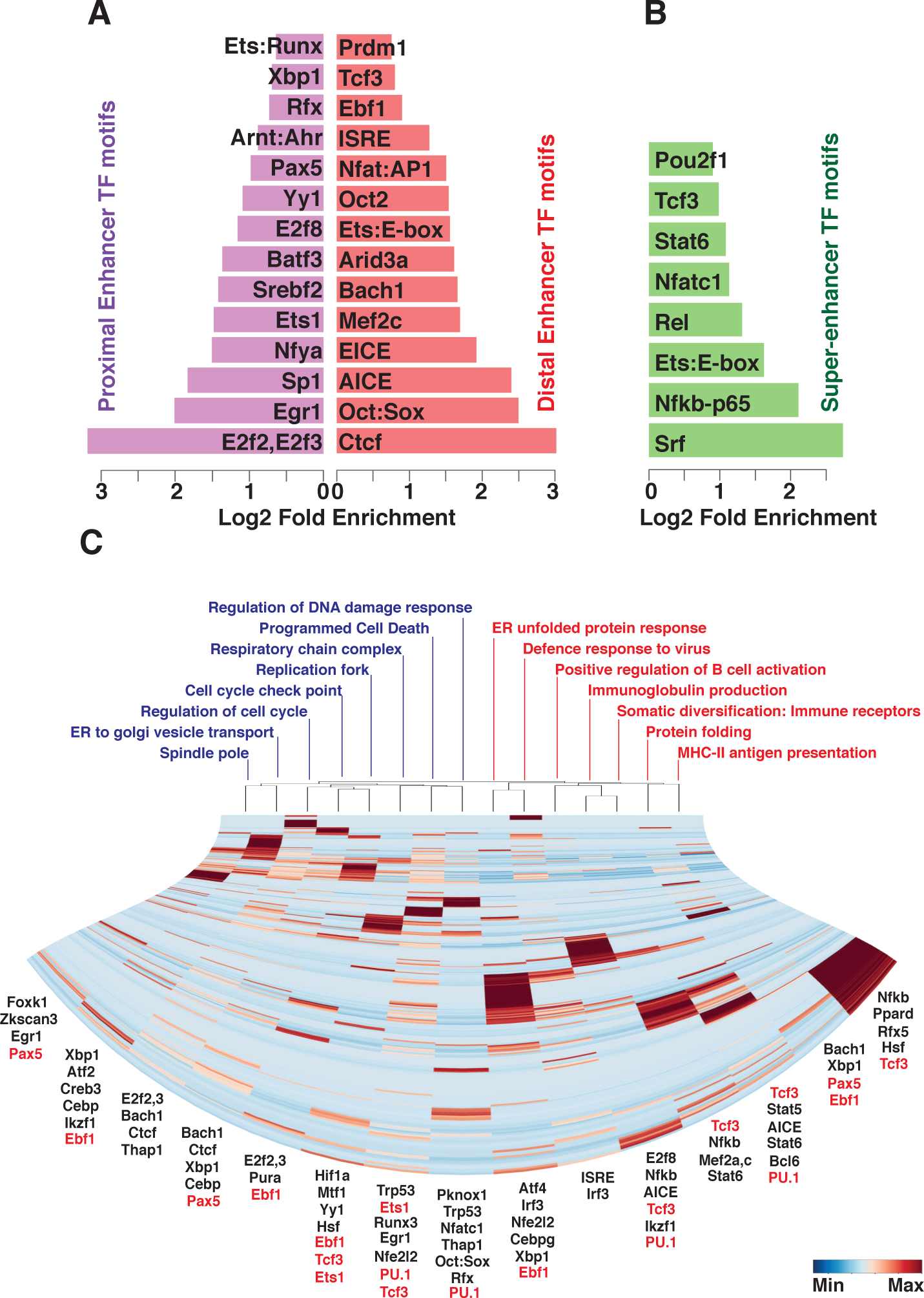
Motif analysis of cis-regulome suggests diverse TF combinatorial codes controlling distinct molecular pathways **A.** TF motif enrichment analysis (HOMER) was performed using STARR-seq tags for promoter proximal versus distal enhancers. TFs whose motifs (simple or composite elements) are highly enriched are displayed in the bar plot with their log2 odds ratios. **B.** TF motif enrichment analysis of super-enhancers as in panel **A**. STARR-seq tags of rest of the cis-regulome (not super-enhancers) was used as the background set. **C.** TF motif enrichment analysis of enhancers interacted with functionally coherent gene modules. Test sets comprised STARR-seq tags overlapping with enhancer co-ordinates which are connected to genes in indicated GO category. Background sets for each GO category comprised of all STARR-seq tags overlapping with enhancers in remaining cis-regulome. Motifs showing significant enrichment (log p-values < −20) were clustered and plotted in heatmap using their log2-fold enrichment values. TF motifs that were not significantly enriched were assigned a value of 0.0. For each GO term the top TF motifs are listed below if they are known or implicated to be involved in regulating that biological process and/or if those TFs are part of that GO term. The TFs highlighted in red font are ones that control B-cell fate determination and/or plasma cell differentiation.

The comprehensive B-cell cis regulome enabled us to analyze large groups of enhancers that were acting on gene sets representing functionally coherent modules. For each comparison the test set contained STARR sequence tags from enhancers connecting to genes in a given pathway, while the background set was sequence tags from rest of the cis-regulome. Clustering was performed using log2 fold enrichment values. To illustrate the utility of this approach, we focused on two types of gene modules (i) those that were reflective of activated, metabolically robust and proliferating cells and (ii) ones that included genes that either are specifically expressed and/or preferentially functioning in differentiating B-cells (Figure 5C). For 15 distinctive GO terms enriched in cis-regulome, a matrix of log2-fold enrichment of TF motifs was generated. This matrix, after clustering, distinguished the two fundamental types of aforementioned gene modules, while simultaneously revealing the TF motif signatures for each molecular pathway. Importantly the TFs acting on these motifs (displayed at the bottom of Figure 5C) were either seen to be part of the GO term that is used to describe the molecular pathway and/or shown to regulate one or more genes in that pathway. This analysis lends support to the predictive power of the cis-regulome. We note that the motifs for the B-lineage-determining TFs (highlighted in Figure 5C) were embedded within the larger and more diverse sets of TF motif combinations. Thus, functional cis-regulome assembly reveals the varied combinatorial TF codes that are likely used to coordinate the expression of gene modules within specific molecular pathways.

### B-cell cis-regulome enriches for combinatorial binding by cell fate determining TFs

To test the relationship between the regulatory sequences that constitute the activated B-cell cis-regulome and the genomic binding landscapes of B-lineage determining transcription factors, we utilized available B-cell ChIP-seq datasets for the following TFs; Ebf1, Pax5, Tcf3 (E2A), PU.1, Ets1 and Prdm1 (Blimp1) (Figure 6A, SRA I.D.s in Methods). Importantly, the binding regions of each of the 6 TFs were more frequently represented in the cis-regulome than in accessible chromatin regions (Figure 6B). To analyze the combinatorial interplay among the 6 TFs, we intersected the ChIP-seq datasets. This revealed that the majority of the binding regions (~70%) involved only one of the six TFs, whereas the remaining regions represented binding peaks for two or more of the six TFs (Figure 6C). As expected for sequence-directed DNA binding, motif enrichment analysis of the genomic regions for DNA sequences recognized by these TFs showed an increase in their frequency with the number of TFs bound per peak (Figure 6D). The foregoing analysis enabled us to probe the relationships between genomic regions that are bound by 1 to as many as 6 of the B-cell TFs and their chromatin accessibility (FAIRE-seq) or enhancer activity (STARR-seq). Chromatin accessibility increased dramatically in frequency when transitioning from genomic regions bound by a single B-cell TF to those bound by 2 or more (Figure 6E). In contrast, the frequency of active enhancers as a function of number of B-cell TFs bound increased more gradually. A genomic region that was bound by 2 of the 6 B-cell TFs had a 50% likelihood of displaying an accessible chromatin structure but a lower probability of functioning as an enhancer. We note that only genomic regions bound by 5 of the 6 B-cell TFs had a 50% likelihood of being active enhancers. This analysis did not preclude the possibility that although the genomic regions being analyzed displayed varying numbers of bound B-cell TFs, their average TF occupancy (including unanalyzed TFs) were comparable. This is highly unlikely as the distribution of the FAIRE-seq tag counts revealed an increase in chromatin accessibility as a function of number of bound B-cell TFs (Figure 6F). Similar analysis of the distribution of STARR-seq tag counts revealed a modest increase in enhancer activity as a function of number of bound B-cell TFs (Figure 6G). Collectively these analyses provide genomic validation for the concept that chromatin accessibility of regulatory elements in the cis-regulome is mediated by a smaller set of TFs, whereas activation of enhancer function requires the interplay of a larger number of TFs. Next, we examined the fraction of the cis-regulome that overlapped with the genomic binding regions of the combinations of the six B-lineage determining TFs. The fraction of the cis-regulome represented by the TF binding regions increased in frequency with the number of TFs bound and displayed a maximum at 4 TFs per peak (Figure 6H). In contrast, accessible chromatin regions, assessed by FAIRE-seq, declined in their overall frequency with increasing number of bound TFs per peak. Therefore, the B-cell cis-regulome, representing a unique subset of all accessible chromatin regions, is strikingly enriched for combinatorial binding regions of B-lineage determining TFs.

**Figure 6.**
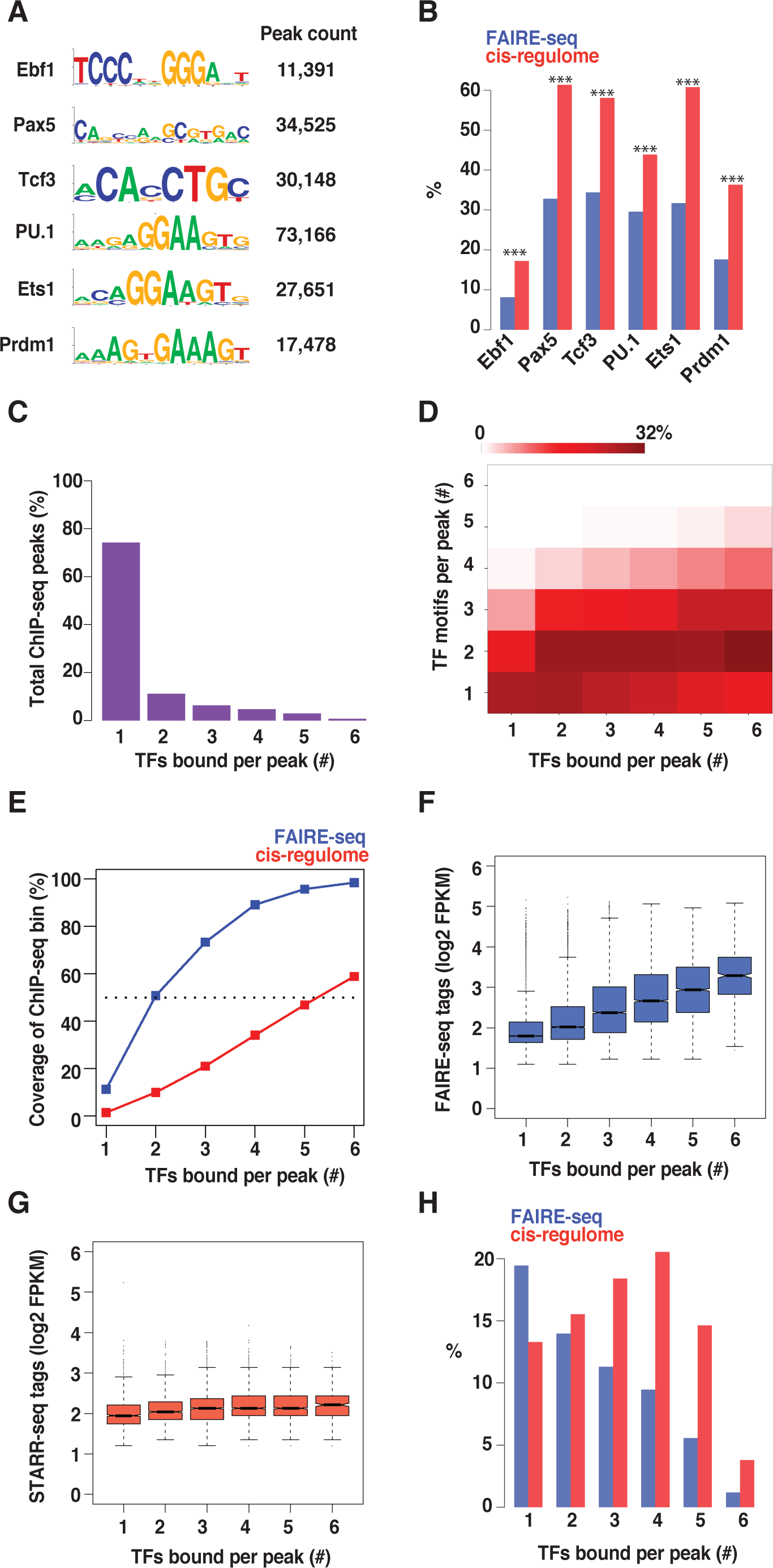
Interaction of B-cell fate determining TFs with the cis-regulome **A.** DNA binding specificities of indicated B-cell fate determining TFs are displayed as sequence logos of their position weight matrices. Published ChIP-seq datasets for these TFs in B-cells were used to determine the counts (indicated) and coordinates of binding peaks for the analyses in subsequent panels. **B**. The fraction of DNA binding peaks for the indicated TFs that overlapped with either the FAIRE-seq peaks (blue) or the B-cell cis-regulome (red) are displayed in bar plots (*** denotes P values < 1E-20, using Fishers exact test) **C.** Combinatorial binding of B-cell TFs in their genomic binding regions. The ChIP-seq peaks for the 6 TFs were binned based on how many represented singular binding events versus those that evidenced binding by 2 or more of the six TFs. **D.** Assessment of TF ChIP-seq bins analyzed in panel **C** for the co-occurrence of TF binding motifs of the same TFs as displayed in panel **D**. Motif co-occurrence frequency (horizontal axis) across each TF ChIP-seq bin (vertical axis) is plotted as a heatmap. **E.** A plot of the fractions of indicated TF ChIP-seq bins (representing binding by one or more of the 6 TFs, analyzed in panel **C**) that reside in accessible chromatin (FAIRE-seq peaks) or are functional enhancers (cis-regulome). **F.** FAIRE-seq peak tag counts for each of the bins analyzed in panel **C**. Boxplots display median values (notches) along with 95% confidence intervals. **G.** STARR-seq peak tag counts for each of the bins as displayed in panel **C**. **H.** Fractions of total FAIRE-seq peaks and cis-regulome enhancers that are covered by TF ChIP-seq bins analyzed in panel **C**.

## Discussion

We have integrated three complementary genomic approaches to assemble and analyze the cis-regulome of a mammalian cell state, namely an activated murine B-cell. The approaches are based on determination of open chromatin regions (structural), high throughput enhancer assays (functional) and delineating enhancer-promoter interactions (connectivity). These three approaches satisfy the hallmark criteria required for the elucidation of a complex mammalian cis-regulome. A notable aspect of our experimental framework is the coupling of the structural approach with one analyzing function i.e., FAIRE-seq with STARR-seq. This enables a facile and directed functional screen of accessible chromatin regions for enhancer activity. The experimental and computational framework is generalizable and will enable elucidation of cis-regulomes for diverse mammalian cellular states.

Although our cis-regulome assembly highlights the integrated functional analysis of presumptive regulatory sequences that are revealed by chromatin profiling, we recognize that it is based on STARR-seq, which has three significant limitations. First of all, STARR-seq utilizes a transient reporter assay which poorly recapitulates chromatin features that may be important for enhancer function (Arnold et al., 2013). Secondly, all enhancers are assayed in the context of a single promoter in the reporter vector. This may result in failure to detect enhancers that show selectivity of action with physiologically relevant promoters (Zabidi et al., 2015). Thirdly, transfection of DNA in mammalian cells activates the interferon response which can bias the reporter assay (Muerdter et al., 2018). We have minimized these limitations by using a large number of independent criteria to validate the enhancers in our B-cell cis-regulome. They are the following: (i) All of the enhancers reside in accessible chromatin regions that are delineated by FAIRE-seq, (ii) vast majority (94%) of the enhancers overlap with DNaseI accessible regions, (iii) large majority (81.3%) of the enhancers are bounded by nucleosomes bearing the activating histone modifications H3K27Ac and/or H3K4me3, (iv) nearly 75% of the enhancers evidence p300 and/or CTCF binding, (v) the enhancers are enriched for eRNAs, (vi) all enhancers are connected to active promoters in B-cells via Hi-C, (Claussnitzer et al.) the enhancers enrich for combinatorial binding by B-cell determining TFs, and (viii) the enhancers enrich for a diverse array of functionally relevant TF motifs and not simply those involved in the interferon response. Thus, this work provides an extensive resource of cross-validated regulatory elements and inferred TFs for future experimental investigations.

We note that STARR-seq identified a significant number of enhancers in the FAIRE-seq library that did not correspond to designated accessible regions. These were precluded from the assembly of the B-cell cis-regulome. Similar findings have been reported with inaccessible chromatin regions in the initial deployment of STARR-seq which involved introduction of a genomic library in Drosophila S2 cells (Arnold et al., 2013). Such regions in both functional screens may represent latent enhancers that are rendered accessible to activating TFs in distinct cellular contexts. However, they could also include DNA fragments with binding sites of one or more TFs that do not function as enhancers in the context of chromatin and thus represent false positives.

The activated B-cell cis-regulome connects 9,989 enhancers (84% of STARR-seq enhancers) to promoters of 7,530 genes (92% of expressed genes). This dense connectivity provides additional support for the biological relevance of the cis-regulome enhancers and also attests to its high degree of completeness for the metabolically activated and proliferating B-cell state. We note that an earlier study reported the assembly of an activated B-cell interactome based on DNaseI-seq, ChIP-seq and ChIA-PET analyses (Kieffer-Kwon et al., 2013). In this case enhancers and promoters were considered to be active based on structural correlates, namely p300, Med12 or Nipbl binding. Furthermore, in this analysis both direct and indirect interactions between promoter and enhancers were included. On this basis, the promoters of 6,890 genes were connected to one or more enhancers. Our approach screens for functional enhancers within accessible chromatin regions and includes only those that are directly connected to active promoters (gene transcript > 4FPKM) via Hi-C. Although our analysis captures several hundred enhancers interacting with plasma cell and germinal center B-cell genes, parallel genomic analyses with these cell populations will be needed to comprehensively assemble their state-specific cis-regulomes.

The activated B-cell cis-regulome is biologically validated by the following considerations. A most compelling finding is that the B-cell cis-regulome not only enriches for the DNA binding motifs of the B-lineage transcriptional determinants PU.1, E2A, Ebf1, Pax5, Ets1 and Blimp1 but displays combinatorial interactions among these TFs (binding peaks for 3 or more) in approximately 60% of the 9,989 enhancers. Furthermore, the super-enhancers are seen to be communicating with genes that are enriched for components encoding the MHCII antigen processing and presentation machinery as well as actin filament branching. These molecular functions are a hallmark of activated B-cells which are programmed to undertake interactions with dendritic cells and cognate T cells in the context of a germinal center response. The super-enhancers are enriched for DNA binding motifs for key signaling-induced TFs such as NF-kB, NFATs, STAT6 and SRF that are known to regulate B-cell activation (Bhattacharyya et al., 2011; Glimcher and Singh, 1999). Based on these findings, we anticipate that hitherto poorly characterized TF motifs that are strongly enriched in the B-cell cis-regulome are likely to be of considerable predictive value. In this regard, we observe enrichment of the Arnt:Ahr composite motif and note that Ahr has recently been shown to regulate B-cell fate choice upon activation (Vaidyanathan et al., 2017). Given that B-cells can be activated to proliferate and differentiate via distinct signaling systems e.g., TLR, BCR, CD40 and cytokines it will be important to determine which enhancers in the LPS induced cis-regulome are shared in other contexts.

Notably, vast majority of genes in the activated B-cell cis-regulome are connected to two or more enhancers with several hundred genes whose promoters were communicating with ten or more enhancers. This cannot be simply explained by the presence of super-enhancers in the cis-regulome as in many cases the multiple enhancers acting on an individual target promoter are not clustered as is the case for the *Prdm1* gene. Although this genomic configuration has also been noted in an earlier study (Kieffer-Kwon et al., 2013) our analysis reveals greater combinatorial diversity and longer distances of action of the distal most enhancers. These findings are intriguing and could simply suggest considerably redundancy in the actions of multiple enhancers on a given promoter, a phenomenon reminiscent of shadow enhancers (Swami, 2010). Alternatively, given that enhancers are considered to contribute to transcriptional bursting (Fukaya et al., 2016), that is reflected by transient and periodic activation of promoters, genes that are acted on simultaneously by multiple enhancers may manifest shorter durations of transcriptional inactivity (Fukaya et al., 2016). In keeping with the latter possibility, genes with higher numbers of active enhancers showed increased levels of nascent transcripts. We note that the B-cell cis-regulome also displays numerous examples of individual enhancers that are communicating with two or more promoters. Furthermore, such enhancers display graded levels of H3K4me1 and H3K27ac in their poised and activated states, respectively, that are correlated with the number of interacting promoters. It remains to be determined if the graded levels of H3K4me1 and H3K27ac displayed by multi-genic enhancers reflect intrinsic differences in their strength of interaction with promoters or are simply a reflection of the number of interacting promoters. We favor the former possibility given that in their poised states multi-genic enhancers are marked with increasing levels of H3K4me1 that are predictive of the number of promoters that they will engage, upon their activation. Regulatory connectivity enabled by multi-genic enhancers could facilitate coordinate control and fine tuning of expression of the target genes in particular signaling contexts. It should be noted that although there are well characterized examples of multiple enhancers acting on a single gene e.g., the immunoglobulin heavy chain locus (Roy et al., 2011) and a single enhancer acting on multiple genes e.g., the IL-4, IL-5 and IL-13 locus (Ansel et al., 2003), it has been unclear if these configurations are the exceptions or the rule for regulatory genome organization. Our analysis of the B-cell cis-regulome reveals that these two organizational features are dominant genome-wide features that generate considerable complexity in enhancer-promoter interactions (see Model, Figure 7).

**Figure 7.**
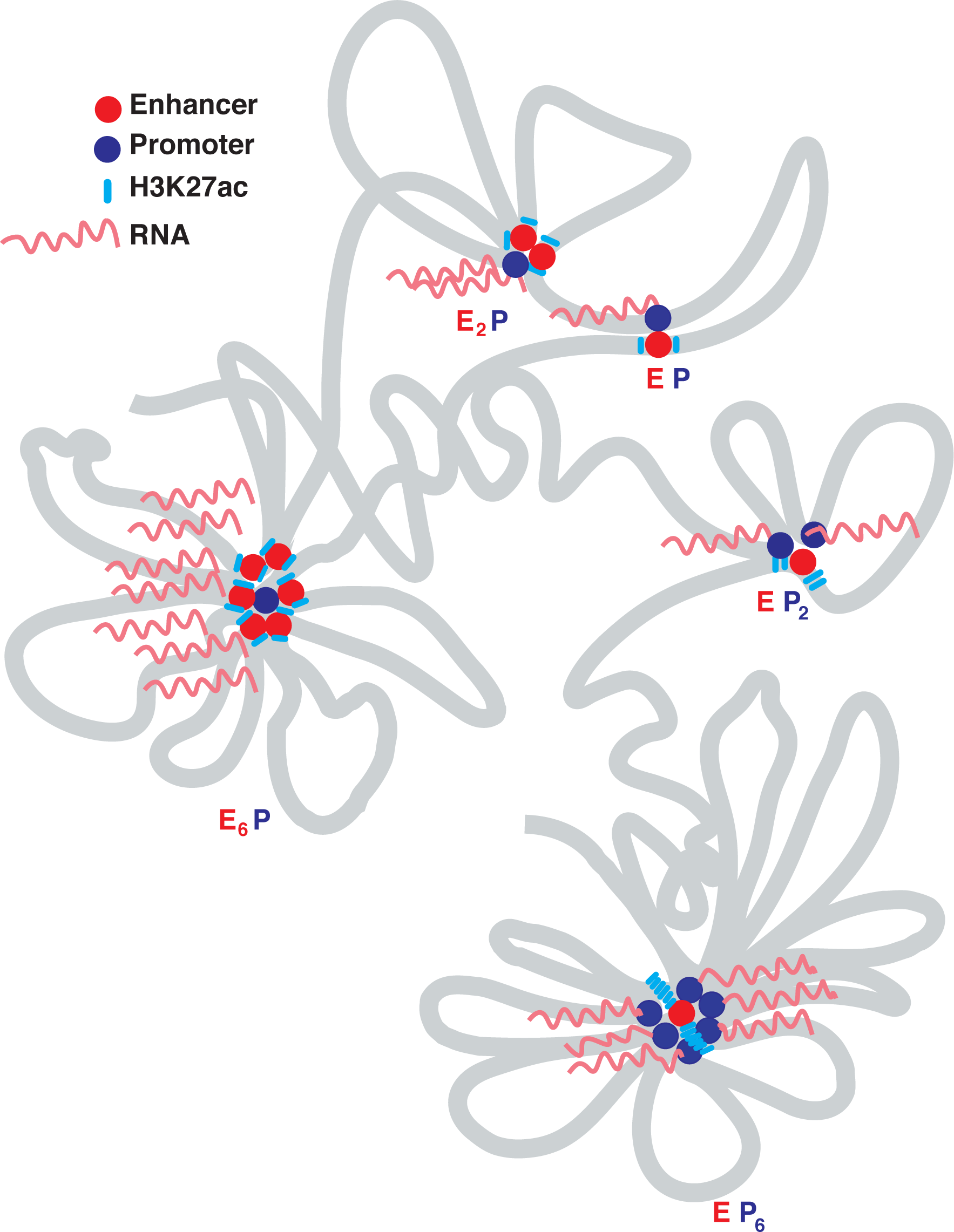
Model of organizational features of cis-regulome The chromosome-based model depicts two dominant cofigurations of the cis-regulome. On the left is shown a promoter (P) of a gene that interacts with 6 distinct enhancers (E_6_P). On the right an enhancer is shown to interact with multiple promoters and genes. The example is of a multi-genic enhancer that can interact with 6 promoters (EP_6_). Although enhancers are displayed as simultaneously interacting with a promoter and vice versa it remains equally possible that the interactions are restricted to individual enhancer-promoter pairs that are temporally distributed.

Another notable feature of the murine B-cell cis-regulome is the very large genomic distances over which enhancers are seen to communicate with target promoters. The median distance for the murine genome is 495kb. Recent evidence has established the ability of enhancers to act over long distances ranging from 0.5 to 1.5Mb (Claussnitzer et al., 2015; Gonen et al., 2018; Johnson et al., 2018). Our global analyses suggest that such long-range action of enhancers is not an exception but rather the rule governing the mammalian regulatory genome. Thus, genomic or locus-specific analyses of the control of mammalian gene activity that focus on regulatory information in proximity to promoters are of limited value given that they ignore regulatory inputs from multiple enhancers that are positioned at very large distances from the promoters.

We propose that the combinatorial complexity of enhancer-promoter configurations (Figure 7) is a manifestation of their rapid evolvability and is made possible by the very large genomic distances (100kb – 10Mb) over which enhancers are seen to communicate with target promoters. Such a large range of enhancer action would enable rapid “enhancer shuffling” via DNA recombination or retro-transposition to generate new alleles of genes. This process of regulatory DNA evolvability would be akin to “exon shuffling” which has been proposed to accelerate intron-mediated evolution of protein coding genes (Long et al., 1995).

Earlier work analyzing the developmental or signaling-induced control of mammalian gene activity has led to the proposal that pioneer TFs are involved in initial alterations of chromatin structure at regulatory sequences that enable accessibility of additional transcription factors leading to activation of enhancer function and gene transcription (Heinz et al., 2010; Mayran and Drouin, 2018; Zaret and Carroll, 2011). Our analyses of the binding landscapes of B-cell determining TFs in relation to accessible chromatin regions and functional enhancers in the cis-regulome provides an important confirmation of this principle on a genome-wide scale. Chromatin accessibility is seen to be associated with the combinatorial interplay of a smaller number of these TFs (2-3) whereas enhancer function appears dependent on a larger set (4-6). This conclusion is reinforced by the enrichment of more diverse combinations of TF motifs that include those for B-cell fate determinants in enhancers connected to functionally coherent gene modules. Notably, our data are not supportive of the concept that individual TFs acting on their own can induce alterations in chromatin structure thereby functioning as “pioneers”. Rather opening of chromatin structure prior to enhancer activation also likely involves combinatorial interplay. This would provide greater specificity of developmentally controlled alterations in chromatin structure mediated by combinatorial interplay of TFs prior to activation of enhancer function.

The delineation of cis-regulomes for distinct mammalian cellular states is essential for the compilation of mammalian regulatory genomes. Cis-regulome assemblies will facilitate the assembly of corresponding complex gene regulatory networks that capture the actions of TFs on the regulatory sequences. Cis-regulomes will need to be continuously refined based on reproducibility and cross-validation analyses and aim for completeness. Thus, it will be necessary to evolve shared experimental and computational frameworks that enable strong collaborative efforts in this arena among scientists focused on particular differentiated cellular states. Our framework is intended to motivate such endeavors.

## Acknowledgements

This study was supported by funds from the Cincinnati Children’s Research Foundation (CCRF) to H.S. We thank core facilities of the Cincinnati Children’s Hospital Medical Center for assistance with mouse colony management, flow cytometry and high-throughput DNA sequencing. We thank Matt Haas for advice with FAIRE-seq and David Fletcher for technical support in sequencing of STARR-seq and eRNA libraries. We are particularly grateful to Lee Grimes, Matt Weirauch, Emily Miraldi and Shekhar Pasare for useful scientific discussions and critical input on the manuscript.

## Materials and methods

### Mice

C56BL/6J mice (6-8 weeks of age) were obtained from the Jackson Laboratory. Mice were housed in specific pathogen-free conditions and used in accordance with guidelines of the Cincinnati Children's Hospital Medical Center Institutional Animal Care and Use Committee.

### B-cell Isolation

Splenic B cells were isolated using B cell isolation kit from Miltenyi (Catalog # 130-090-862) and cultured in complete RPMI with 10% FCS. Purified B cells (250,000 cells/ml) were activated with LPS (10μg/ml, *Salmonella typhimurium*, Sigma Catalog # L6143).

### RNA-seq

RNA isolation and cDNA library preparation were performed as described previously (Xu et al., 2015). Paired end sequencing was conducted on Illumina Hi-seq 2500 platform.

### Formaldehyde crosslinking of B-cells

Crosslinking of resting and LPS activated B-cells was performed with aliquots of 5 x 10^6^ cells in 1 ml of 1% formaldehyde, 10 mM NaCl, 0.5mM EGTA, 0.1mM EDTA and 5mM HEPES pH 8.0, for 10 minutes at room temperature (RT). Crosslinking reactions were quenched with 55 μl of 5M glycine for 5 minutes at RT and cells were pelleted at 2000g for 5 min. After washing with ice cold PBS, cell pellets were stored at −80°C. Cell pellets were used for FAIRE-seq, ChIP-seq, and Hi-C analyses.

### Histone ChIP-seq

ChIP-seq was performed with the following antibodies; rabbit polyclonal anti-H3K27ac antibody (Cat. No. C15410196, Diagenode Inc.), rabbit monoclonal anti-H3K4me3 antibody (Cat. No. 17-614, Millipore) and rabbit polyclonal anti-H3K4me1 antibody (Cat. No. C15410194, Diagenode Inc.) as described previously (Rochman et. al. 2018).

### FAIRE-seq

Frozen, crosslinked B-cell pellets (5 x 10^6^ cells) were thawed on ice and resuspended in I ml of L1 lysis buffer (140mM NaCl, 1mM EDTA, 10% glycerol, 0.5% NP40, 0.25% Triton X-100, 50mM HEPES pH 7.5 and protease inhibitors) and incubated at 4°C for 30 minutes with rotation. The cellular lysate was centrifuged (1,000g for 5 minutes) and the chromatin containing pellet was resuspended in 1 ml of L2 buffer (200mM NaCl, 1mM EDTA, 0.5mM EGTA, 10mM Tris-HCl pH 8.0 and protease inhibitors) and incubated at 4°C 10 minutes with rotation. After centrifugation as above the chromatin containing pellet was resuspended in 1 ml sonication buffer (1% Triton X-100, 0.1% deoxycholate, 5mM EDTA, 150mM, NaCl, 50mM Tris-HCl pH 8.0) and sonicated in a Covaris instrument (10% duty cycle, 175 pulse and 200 burst, 7 minutes). Sonicated chromatin suspension was centrifuged at 17,000g for 10 minutes and the supernatant was subjected to Phenol:Chloroform:Isoamyl alcohol (25:24:1) extraction. The FAIRE DNA was recovered from the aqueous phase by ethanol precipitation, quantified using NanoDrop and then used to generate FAIRE-seq library. Adapters were ligated to FAIRE DNA (300 ng) using NEBNext^®^ Ultra™ II DNA Library Prep Kit (NEB Catalog No. E7645S). After U excision step, one-third of the DNA sample was amplified using Illumina indexing primers (NEB Catalog no. 7600L). PCR product was size selected by running on 1.5% agarose gel. DNA fragments ranging in size from 300-700 bp were purified and sequenced on Illumina HiSeq 2500 to generate 75 bp paired-end reads.

### STARR-seq

STARR-seq libraries were generated from remaining two-thirds aliquot of FAIRE-DNA that had undergone adapter ligation and U excision (see FAIRE-seq protocol). PCR (4 reactions) was performed using the following STARR-seq primers (fwd: TAG AGC ATG CAC CGG ACA CTC TTT CCC TAC ACG ACG CTC TTC CGA TCT, and rev: GGC CGA ATT CGT CGA GTG ACT GGA GTT CAG ACG TGT GCT CTT CCG ATC T) and amplification conditions (98°C for 90 seconds followed by 12 cycles of 98°C for 15s, 65°C for 30s, 72°C for 30s) (Arnold et al., 2013). Pooled amplified product was size-selected (300-700 bp) by resolving on 1.5% agarose gel. Size-selected DNA fragments were used for cloning into STARR-seq vector, linearized with AgeI and SalI. Directional cloning into STARR-seq vector was performed with Takara In-Fusion HD kit in 10 reactions with 1:3 molar ratio of linearized-plasmid to FAIRE-fragments. Reactions were pooled, and after ethanol precipitation DNA was then electroporated into MegaX DH10B™ T1R Electrocomp™ Cells in 6 reactions. After incubation in recovery medium at 37°C for 1 hour, cell suspensions were pooled and grown in 500ml LB with carbenicillin (100 μg/ml) for 12-16 hours (OD_600_ = 0.8 - 1). We note that each transformation reaction yielded up to 10^9^ CFU. Plasmid DNA (STARR-seq library) was purified using QIAGEN columns. Inserts in STARR-seq library were sequenced using a two-step, nested PCR strategy. First step involved 6 cycles of amplification with the following primers (fwd: GGGCCAGCTGTTGGGGT G*A*G*T*A*C, rev: CTTATCATGTCTGCTCG A*A*G*C, where * indicate phosphorothioate bonds) (Vanhille et al., 2015) (98°C for 5min followed by 6 cycles of 98°C for 15s, 65°C for 30s, 72°C for 70s). DNA from multiple reactions was pooled, concentrated using Ampure-XP beads and inserts were purified away from template plasmid DNA by agarose gel electrophoresis. Resulting product was then used for second-step PCR (98°C for 90 seconds followed by 10 cycles of 98°C for 15s, 65°C for 30s, 72°C for 30s) using Illumina indexing primers (NEB Catalog No. 7500L). DNA was purified using Ampure XP beads and sequenced on Illumina HiSeq 2500 to generate 75 bp paired-end reads. Sequencing of the STARR-seq library revealed >81% coverage of the input FAIRE-DNA.

For each STARR-seq experiment, 6-8 transfections were performed using aliquots of 5 million activated B cells and 15 μg of STARR-seq plasmid library. LPS activated (48h or 60h) splenic B-cells were transfected using Neon electroporation kit with R buffer and Neon Transfection System (1350V, single 40ms pulse). B-cells were washed with ice cold PBS before resuspending in R buffer. After electroporation, B-cells were immediately re-plated in 3 ml of complete medium with 10 μg /ml LPS for 24 or 12 hours depending on the timepoint of transfection. At 72h post LPS activation, B cells from an individual experiment were pooled and used to isolate RNA with RNeasy Mini kit (QIAGEN). After treatment with Ambion Turbo DNase, the RNA was purified using RNeasy MinElute kit (QIAGEN). mRNA was isolated using Oligo-dT beads and first strand synthesized using STARR-seq cDNA primer (CAA ACT CAT CAA TGT ATC TTA TCA TG). All subsequent steps of STARR-seq protocol were as described previously (Arnold et al., 2013). DNA sequencing was performed on Illumina HiSeq 2500 platform to generate 75 bp paired-end reads.

### Luciferase Reporter Assay

STARR-seq regions listed below were amplified using genomic DNA and the indicated primer pairs. Amplified DNA fragments were cloned into pGL4.26 plasmid (Promega Catalog No. E8441). All constructs were sequenced to verify their identity. Luciferase assays were performed in CH12 cells by co-transfecting 10 μg of with test plasmid and 0.5 μg of TK-Renilla luciferase reporter plasmid (Promega Catalog No. E6931) as control. Transfections were performed Neon electroporation kit with R buffer and Neon Transfection System (1350V, single 40ms pulse). CH12 cells were washed with ice cold PBS before resuspending in R buffer. After electroporation, the cells were immediately re-plated in 2 ml of complete medium for 24 hours and then lysates were analyzed with the Dual Luciferase reporter system (Promega Catalog No. E1910).

### Hi-C

*In-situ* Hi-C was performed on cross-linked LPS activated (72h) B-cells as described by (Rao et al., 2014) with some minor modifications. Aliquots of 5 million cross-linked B-cells were used to prepare nuclei, that were subjected to 3 hours of digestion with MboI followed by end filling to create biotin labeled blunt ends. Blunt ends were ligated using T4 DNA ligase at 16°C for 2 hours. After ligation, nuclei were digested with Proteinase K and genomic DNA was isolated using phenol:chloroform:isoamyl alcohol (25:24:1) followed by ethanol precipitation. DNA was sonicated using Covaris instrument (Duty cycle: 15, PIP: 500, Cycles/Burst: 200 and Time: 58 seconds) to obtain fragments ranging in size from 200-800 bp. Biotinylated DNA was purified using Pierce Streptavidin magnetic beads (Thermo Fisher Catalog No. 88816). While on beads, DNA ends were blunt ended and dA tailed as described in (Rao *et. al.*), followed by ligation of Illumina sequencing adapter with T4 DNA ligase. After U excision step, DNA samples were amplified using Illumina indexing primers (NEB Catalog no. 7600L). PCR reactions were performed in 16 separate 20 μl aliquots on beads involving 8 cycles of amplification (98°C for 90 seconds followed by 98°C for 15s, 65°C for 30s, 72°C for 30s). PCR product was separated using Dynamag magnetic separator. Supernatants were pooled and concentrated using AMPureXP beads ratio 1.8:1. Concentrated DNA library was size selected using 1.5% agarose gel to obtain fragments ranging in size from 400-700 bp. Sequencing was performed on Illumina Hi-Seq 2500 to obtain 75 bp paired end reads. For each biological replicate, ~300 million reads were generated.

### Nascent RNA-seq

LPS activated B-cells (5 million) were pulse labeled with EU (2.5 mM, Invitrogen Catalog No. C10365) for 30 minutes prior to harvesting at 72h post activation. Cells were pelleted and snap frozen using dry ice. To prepare RNA, cells were thawed on ice for 15 minutes and then nuclei were prepared by resuspending cells in NIB-250 (15mM Tris-HCl pH 7.5, 60mM KCl, 15mM NaCl, 5mM MgCl_2_, 1mM CaCl_2_, 250mM Sucrose and protease inhibitors) and 0.3% NP-40 on ice for 5 minutes. Nuclei were pelleted by centrifugation at 600g for 5 minutes at 4^0^ C. Nuclear pellets were washed twice with NIB-250 and then lysed with 1 ml of Trizol. RNA was isolated manufacturer’s protocol and quantified. Nuclear RNA (2 ug) was depleted of rRNA using NEB kit Catalog No. E6310S). mRNA was removed by incubation with oligo-dT magnetic beads for 10 minutes. Supernatant was aspirated using Dynamag magnetic separator and RNA was precipitated using ethanol. Sample was subjected to Click-IT chemistry to enable biotinylation of EU present in nascent transcripts. Biotin pull down of EU-labeled nascent RNA and all subsequent steps were performed as per Invitrogen kit instructions. Superscript VILO (Invitrogen Catalog No. 11756050) was used to synthesize cDNA on beads. After second strand synthesis the double stranded DNA was purified using AmpureXP beads at 1.8:1 ratio. Resulting DNA (~0.4 ng) was used to make sequencing library with Thruplex DNA-seq kit (Rubicon Genomics, Catalog. No. R400523). Sequencing was performed on Illumina Hi-Seq 2500 to obtain 75 bp paired end reads.

### Data processing and computational analysis

#### RNAseq

Reads were aligned using Tophat2 (Kim et al., 2013) with options “— no-disconcordant –no-mixed –maxhits 1” to mm9 genome. Cufflink was used to determine transcript abundance in FPKM. Genes whose transcripts were detected at a value of 4 FPKM or higher were used in the assembly of the cis-regulome.

#### Nascent RNA-seq

Reads were aligned using STAR-aligner (Dobin et al., 2013) with default options and output files were then processed using Homer suite of tools (Heinz et al., 2010). The Homer function “analyzeRepeats.pl” was used to calculate read counts on either enhancers or gene bodies of expressed genes. Only unique reads were counted.

#### Histone ChIP-seq analysis

H3K27ac, H3K4me3 and H3K4me1 ChIP-seq reads were aligned to mm9 genome with default bowtie2 options (Langmead and Salzberg, 2012). Unique reads were used for peak calling with Homer function findPeaks with a 500 bp sliding window using option “-style histone”. For calling super-enhancers we used option “-style super” with default settings. Peaks from the blacklisted regions of mm9 genome were discarded. The function “analyzeRepeats.pl” to determine quantile normalized tag counts for ChIP-seq peaks.

#### FAIRE-seq and STARR-Seq

Paired end reads were aligned using bowtie options “—very-sensitive –k 1 –m 500” and output sam files were converted to bam files. Then bamToBed tools were used to convert bam files into paired end bed files in “bedpe” format. A custom R script was used to filter out spurious alignments such as reads aligned to random scaffold, reads mapping to mitochondrial DNA, or paired reads with ends mapping to different chromosomes. Tags were generated using start and end coordinates derived from paired alignment of forward and reverse reads, and only unique tags were considered in the following analysis. Tags longer than 1kb were discarded. Bed files containing tag information were analysed with Homer suite of tools. Peak calling was done using function “findPeaks” with options “-size 300 –minDist 300 –L 0 –center-fdr 0.05”. Peaks were 300bp in size and centered on maximum tag density. Peaks were separated by at least 300bp. For peak calling in STARR-seq the tags from replicate experiments were combined. Peaks from blacklisted regions in mm9 genome were discarded.

#### Hi-C

Hi-C paired end reads were aligned separately with default bowtie options to mm9 and up to 2 best alignments were kept considering that a chimeric junction can reside in one of the paired ends. Alignment output was used as input data for Homer functions (makeTagDirectory) to generate primary Hi-C tag directory. Tags which were result of self-ligation, ends close to a “GATC” site in mm9 genome, were discarded. Tags where read ends were closer than 900 bp, were eliminated. Background models were generated for each bin size using function “analyzeHiC –bgonly”. Next Homer function “findHiCInteractionsByChr” was used to selectively analyze cis-regulatory interactions only, for bin sizes of 1kb, 2kb, 5kb and 10kb with overlapping regions of varying sizes to avoid penalizing elements at the boundaries of bins. For bin resolution of 5 kb, sliding windows of 1 kb and 2.5 kb were used and for bin resolution of 10 kb, sliding windows of 2.5 kb and 5 kb were used. 1kb and 2kb bin resolutions used 500bp and 1kb sliding windows, respectively. Interactions with at least 5 interaction read pairs with p-values less than 0.0001 were used in assembly of cis-regulome. For plotting the Hi-C contact distances we used Homer assigned unique interaction ids.

#### Cis-regulome assembly

Analysis of the FAIRE-seq and STARR-seq datasets generated 11,809 overlapping peaks. A minimal overlap between a FAIRE-seq and STARR-seq peak of a single bp was allowed in the analysis. We note that a window size of 300bp is small enough for a shift in the density of tags based on structural (FAIRE-seq) versus functional (STARR-seq) features. Thus centering of the two types of peaks based on highest tag density is not expected to generate complete peak alignment. Importantly, 93% of these peaks had overlaps of 100 bp or more. These 11,809 peak coordinates along with those of 8,215 promoters (−1000bp, TSS, +100bp) representing expressed genes in activated B cells were queried for Hi-C interactions using 4 bin sizes (10, 5, 2 and 1kb). A custom R script to was used to identify 2 interacting bins such that one bin overlapped with one or more enhancers while the other overlapped with at least one promoter of an expressed gene. This analysis yielded the activated B-cell cis-regulome comprising of 9989 enhancers connected to 7530 expressed genes. We note that only direct Hi-C contacts for enhancer promoter pairs were used for cis-regulome assembly. For example, in the context of the following interactions: enhancer1-enhancer2-promoter1-promoter2-enhancer3 only enhancer2-promoter1 and promoter2-enhancer3 would be considered for the cis-regulome build.

#### Transcription factor ChIP-seq and DNaseI–seq analysis of public datasets

Published transcription factor ChIP-seq and DNaseI-seq data sets were obtained from the SRA archive (SRA ids provided below) and aligned to mm9 genome with default bowtie2 options. We merged all replicates for any given TF ChIP-seq and performed the peak calls using Homer function findPeaks with following options {-style factor -size 300-minDist 300 -center -tbp 1}. To count binding of one or more TFs in peak regions, overlaps of peaks were determined in R using function subsetByOverlaps from “GenomicRanges” package. Peaks were called in DNaseI-seq data using following options {-size 150 -minDist 150 -center -L 0-tbp 1 -style dnase}.

#### Gene Ontology analysis

Gene ontology (GO) based enrichment of gene sets associated with super-enhancers was performed using conditional GO testing from GOstats package (Falcon and Gentleman, 2007).

#### TF-motif analysis

A custom TF motif library was compiled (Supplementary Data S4) for all TFs expressed in activated B cells (RNA-seq dataset) using cis-DB (Weirauch et al., 2014). This compilation was supplemented with TFs which are known regulators of plasma cell or GC B cell differentiation. All PWMs were trimmed for edges with lower information content (<0.5). A similarity matrix for all PWMs was generated using Homer function “compareMotifs.pl” with option “-matrix”. For any TF with multiple PWMs a minimal spanning tree, with the PWMs as nodes, was generated using the similarity matrix. The PWM with most connecting edges was selected and the others discarded. In the case of ties, the PWM with highest information content was chosen. In cases where 2 or more TFs had PWMs with similarity score of more than 0.9 then were all represented by a single PWM with the highest information content. Gene symbols of these TFs separated by “:” were used as a naming convention in the final library. PWM cut off scores were assigned based on information content for a given PWM. For PWMs with maximal scores (Function maxScore from Biostrings) of >10, 6-10 and <6, cutoffs of 85%, 90% and 95% respectively, were used. All PWM filtering was performed in R with custom written scripts. This library was written in minimal Homer format for PWMs. Homer vertebrates PWM library, which contains ChIP-seq derived motifs including composite elements was added to the above compilation to generate final library. No trimming, filtering or score adjustments were performed on TF motifs in Homer vertebrates PWM library. TF motif enrichment was performed either using coordinates of STARR-seq peaks or those of STARR-seq tags that overlapped with STARR-seq peaks. The latter was advantageous from the standpoint of functional enrichment and afforded greater statistical power given that a minimum of 10-15 tags were associated with a given peak.

#### Genome Tracks

For genome tracks the Homer tag directories of H3K27ac ChIP-seq, FAIRE-seq and STARR-seq were used to generate the track files using function makeUCSCfile. Output files were then imported in R and “Sushi” (Phanstiel et al., 2014) was used to generate the genome track plots. All genomic overlaps were determined in R using package “GenomicRanges” (Lawrence et al., 2013).

#### Graphical representations and Statistical tests

All graphical plots were created in R with built in functions along with add in routines from ggplot2. All statistical analyses were performed in R with the exception where P values are reported using Homer.

---

Public data set used in study.

**Public data set used in study:**
|  |  |  |  |
| --- | --- | --- | --- |
| SRR409118 | Ebf1 | SRR974813 | DNaseI aB |
| SRR499700 | Pax5 | SRR974814 | DNaseI aB |
| SRR499701 | Pax5 | SRR974816 | DNaseI aB |
| SRR499702 | Pax5 |  |  |
| SRR499703 | Pax5 |  |  |
| SRR499704 | Pax5 |  |  |
| SRR499705 | Pax5 |  |  |
| SRR499706 | Pax5 |  |  |
| SRR3158144 | E2A |  |  |
| SRR3724509 | Ets1 |  |  |
| SRR3724510 | Ets1 |  |  |
| SRR372712 | Ctcf |  |  |
| SRR372713 | Ctcf |  |  |
| SRR1014532 | PU1 |  |  |
| SRR1003255 | P300 |  |  |
| SRR5087645 | Prdm1 |  |  |
| SRR5087646 | Prdm1 |  |  |
| SRR5087647 | Prdm1 |  |  |
| SRR5087648 | Prdm1 |  |  |
| SRR5087649 | Prdm1 |  |  |
| SRR5087650 | Prdm1 |  |  |
| SRR5087651 | Prdm1 |  |  |
| SRR5087652 | Prdm1 |  |  |
| SRR5087653 | Prdm1 |  |  |
| SRR5087654 | Prdm1 |  |  |
| SRR5087655 | Prdm1 |  |  |
| SRR5087656 | Prdm1 |  |  |
| SRR5087657 | Prdm1 |  |  |
| SRR5087658 | Prdm1 |  |  |
| SRR5087659 | Prdm1 |  |  |
| SRR5087660 | Prdm1 |  |  |

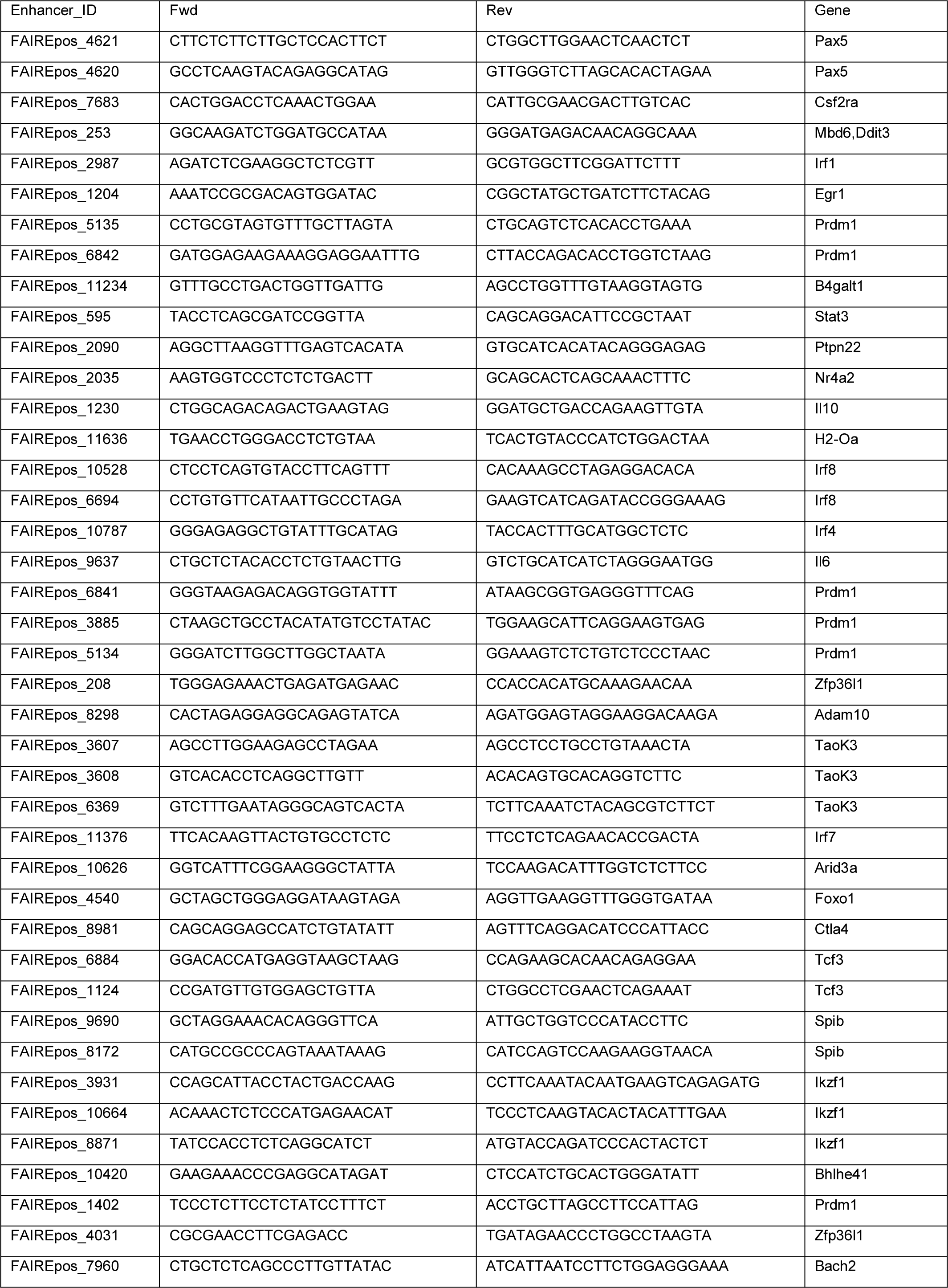
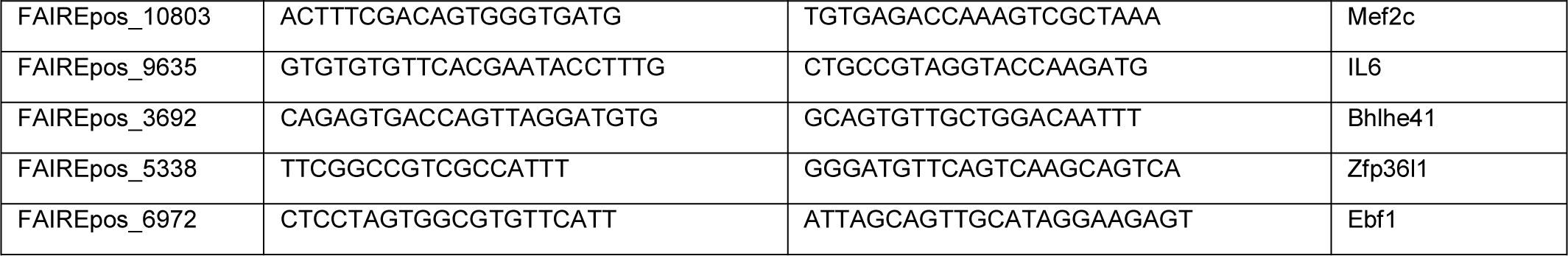
Enhancer Luciferase Reporter Constructs.

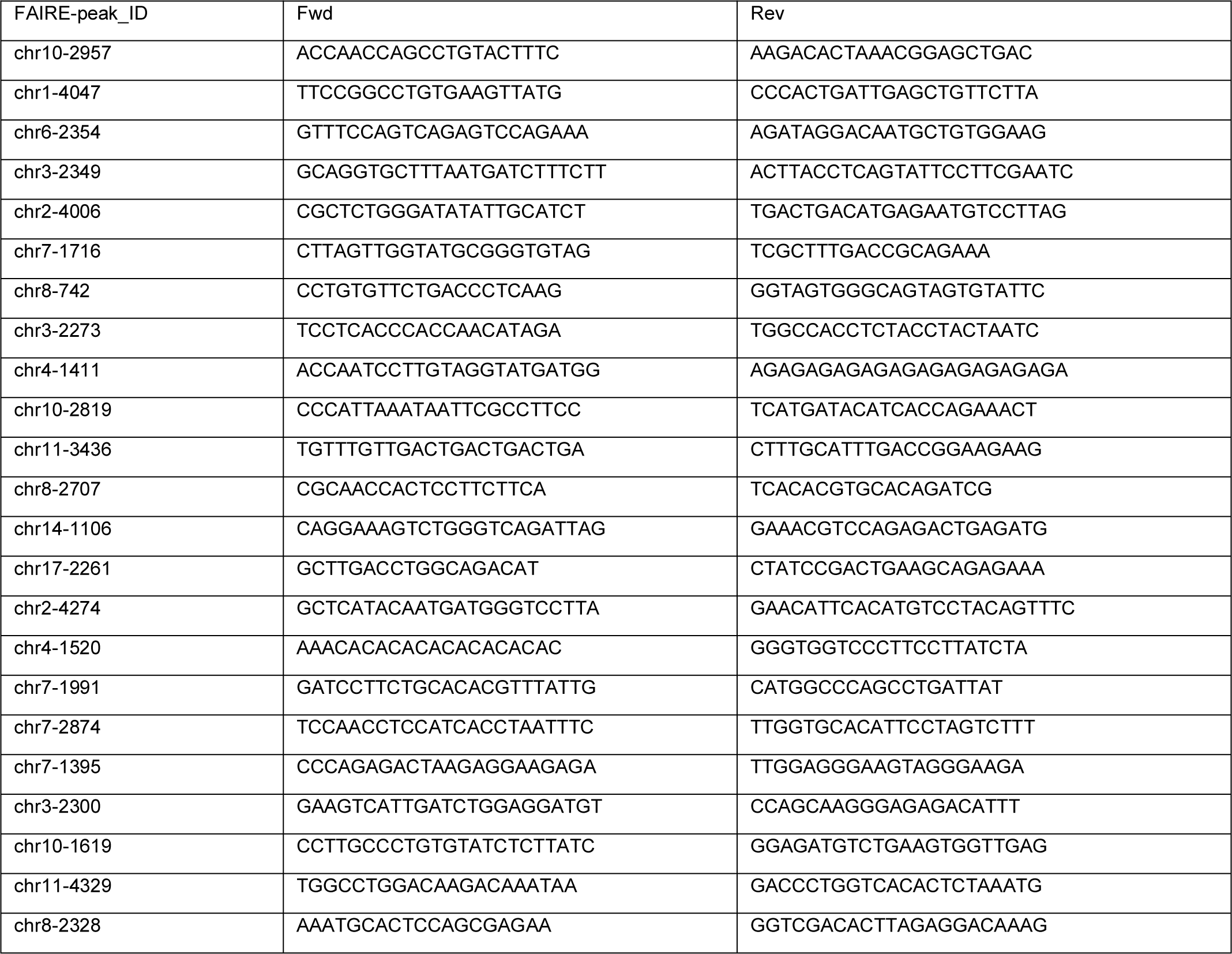
Control Luciferase Reporter Constructs.

**Computational packages:**

- R version 3.3.0 (2016-05-03)
  - Sushi_1.10.0
  - biomaRt_2.28.0
  - GOstats_2.38.1
  - Matrix_1.2-12
  - AnnotationDbi_1.34.4
  - Biobase_2.32.0
  - seqinr_3.4-5
  - rtracklayer_1.32.2
  - GenomicRanges_1.24.3
  - GenomeInfoDb_1.8.7
  - IRanges_2.6.1
  - S4Vectors_0.10.3
  - BiocGenerics_0.18.0
  - Biostrings_2.40.2
  - GenomicAlignments_1.8.4
  - GO.db_3.3.0
  - org.Mm.eg.db_3.3.0
  - AnnotationForge_1.14.2
  - ade4_1.7-10
  - genefilter_1.54.2
- Bowtie2 (DNA reads aligner) version 2.2.3
- Tophat2 (RNA reads aligner) version 2.0.13
- Homer (analysis suite for mapped read data) version 4.9
- STAR (RNA reads aligner) version

**Figure S1.**
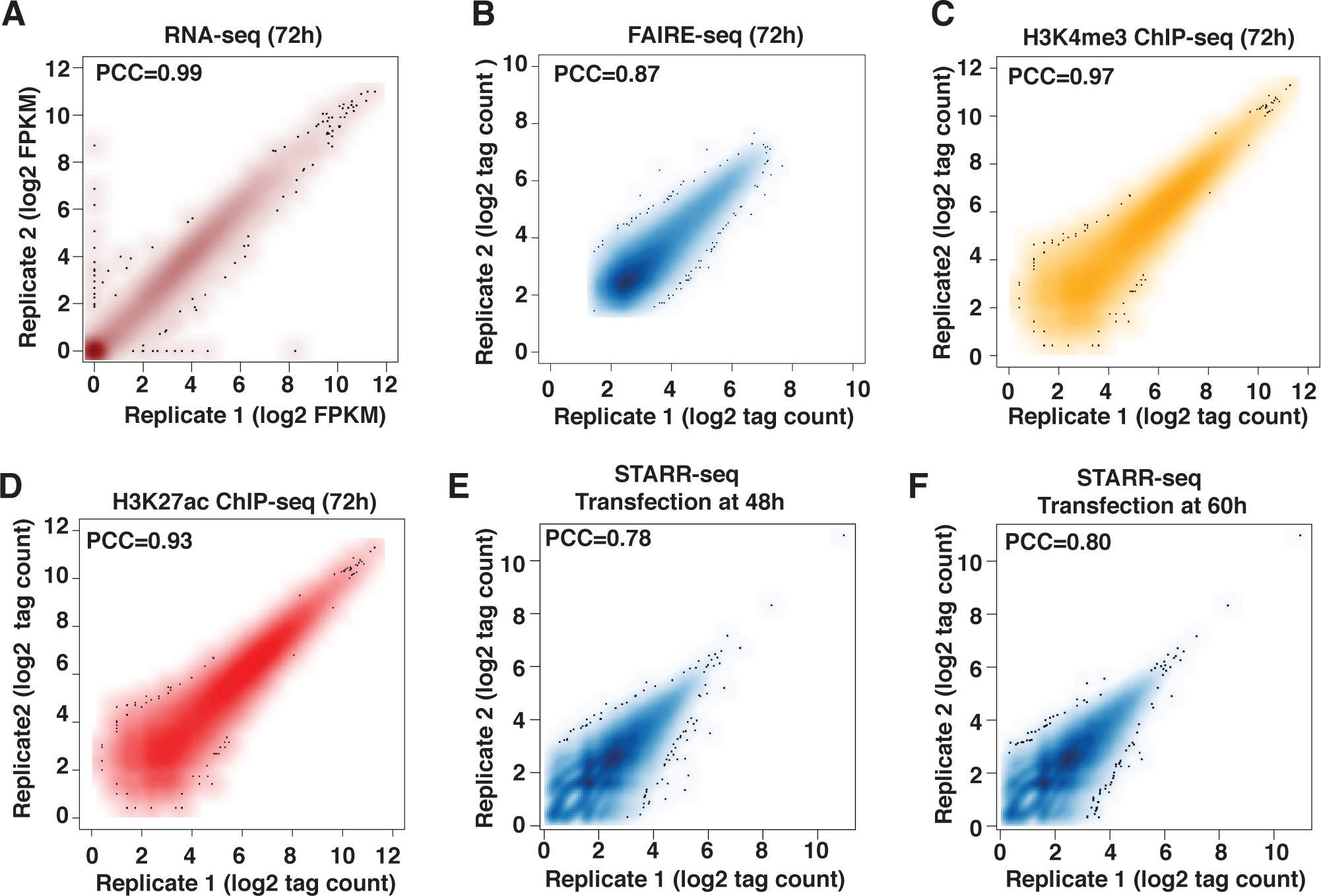
Reproducibility of RNA-seq, chromatin profiling and STARR-seq in LPS activated B-cells. Analyzed data is from biological replicates. **A.** RNA-seq data is plotted for all known genes as log transform of FPKM values. **B.** FAIRE-seq peaks are plotted as quantile normalized tag counts for 55,130 regions **C, D.** H3K27ac peaks and H3K4me3 peaks plotted for quantile normalized tag counts, respectively. **E, F.** A pool of size-selected FAIRE-DNA fragments were used to construct two libraries, each of which was transfected into LPS activated splenic B-cells at either 48h or 60h. RNA was isolated at 72h post-activation for sequencing. Quantile normalized tag counts of STARR-seq output for the 55,130 FAIRE-seq peaks are plotted. In all panels, a value of 1 was added to the counts before transforming to log2. Indicated PCC values are Pearson correlation coefficients for biological replicates.

**Figure S2.**
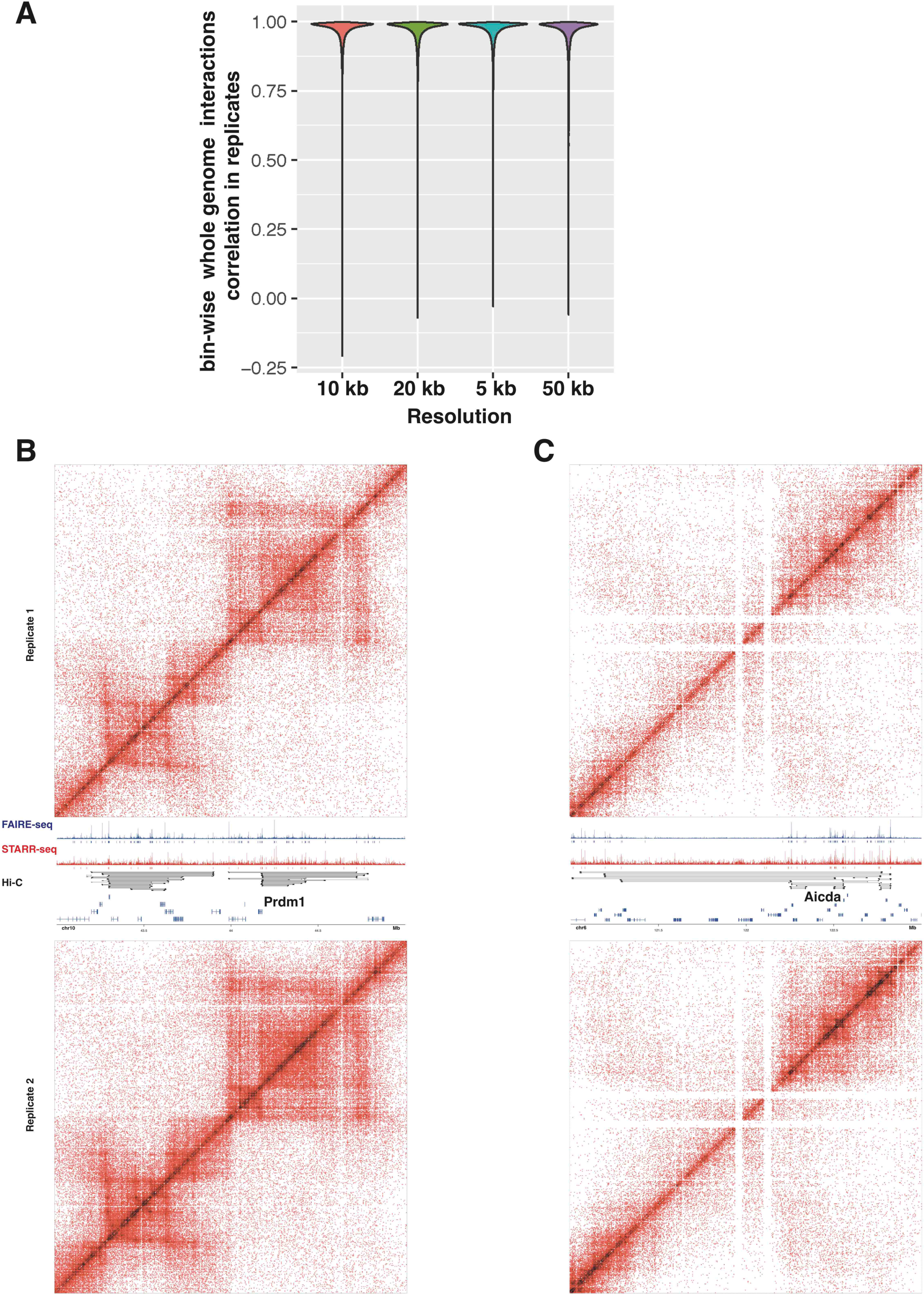
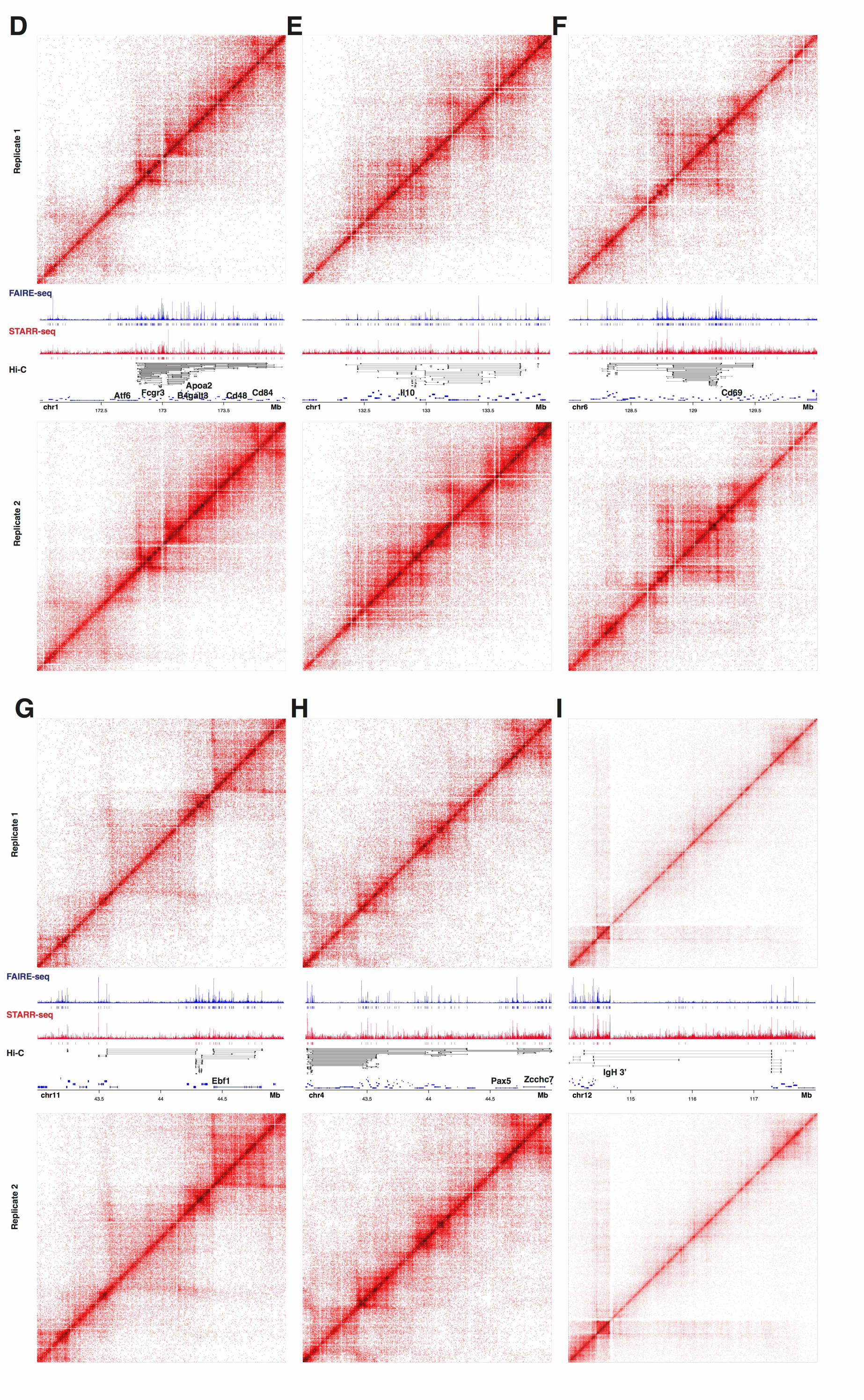
Reproducibility of Hi-C analyses. **A.** Hi-C replicates of LPS activated B-cells (72h). Biologically independent experiments were analyzed for reproducibility by bin-wise genome wide correlation for the indicated bin-sizes. The Homer function getHiCcorrDiff.pl was used to generate correlation matrices. Correlation coefficients of each bin at a given resolution are plotted. **B-I.** High resolution contact matrices (5 kb bin size), using raw counts for the indicated biologically important loci, are displayed (replicates). FAIRE-seq (blue) and STARR-seq (red) tracks are displayed below the contact matrix with their peak calls. All enhancer-promoter connectivity links for indicated locus in the combined Hi-C dataset (replicate 1 and 2) are shown with black bars below the STARR-seq tracks.

**Figure S3.**
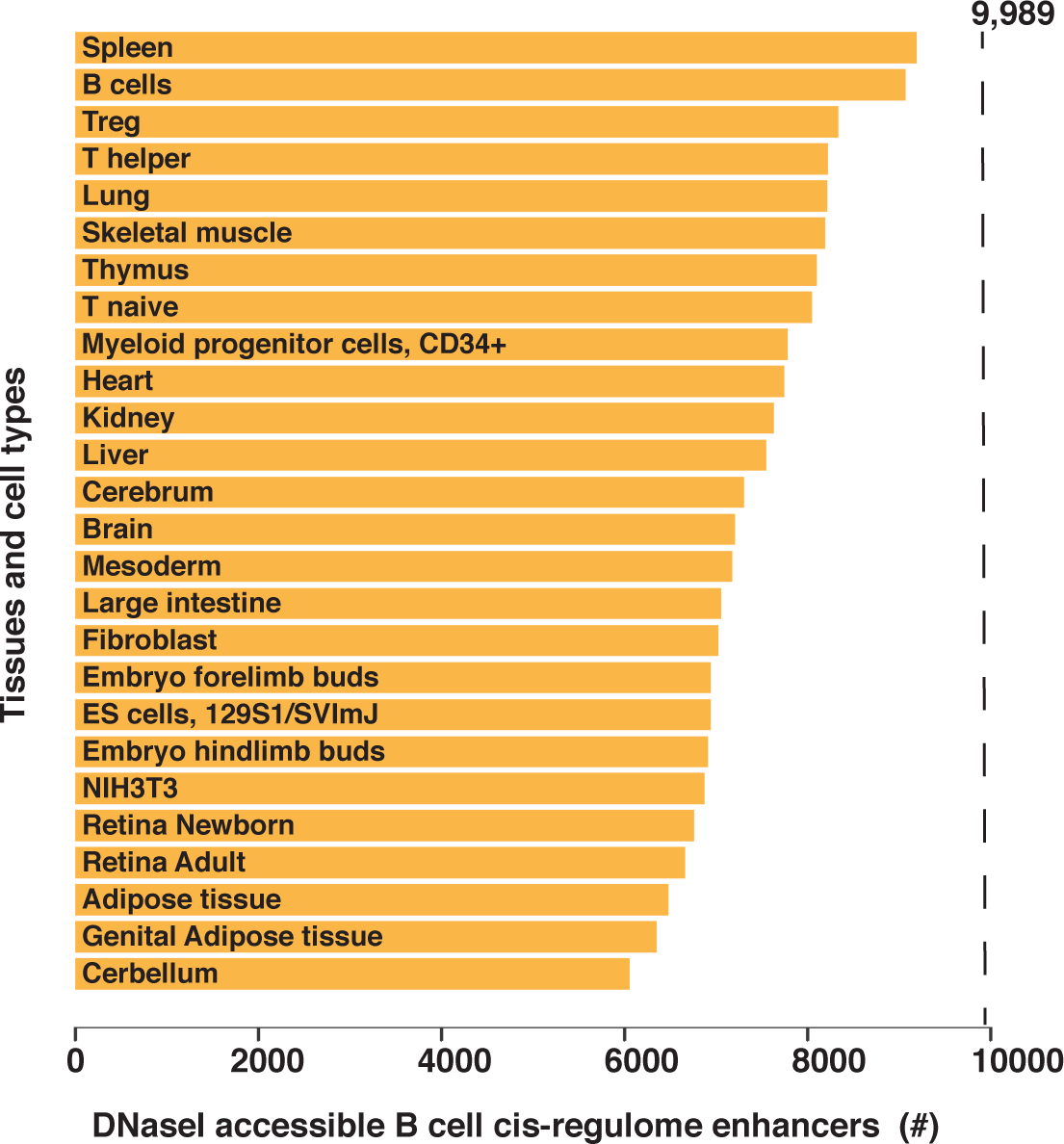
DNaseI accessibility of B-cell cis-regulome enhancers in different cell types and tissues. Mouse ENCODE data for DNaseI accessible regions was obtained from UCSC (genome.ucsc.edu/encode/downloadsMouse.html) in narrowpeak file formats. Counts of cis-regulome enhancers (total of 9,989) overlapping with DNaseI accessible regions of indicated cell and tissue types are plotted here.

**Figure S4.**
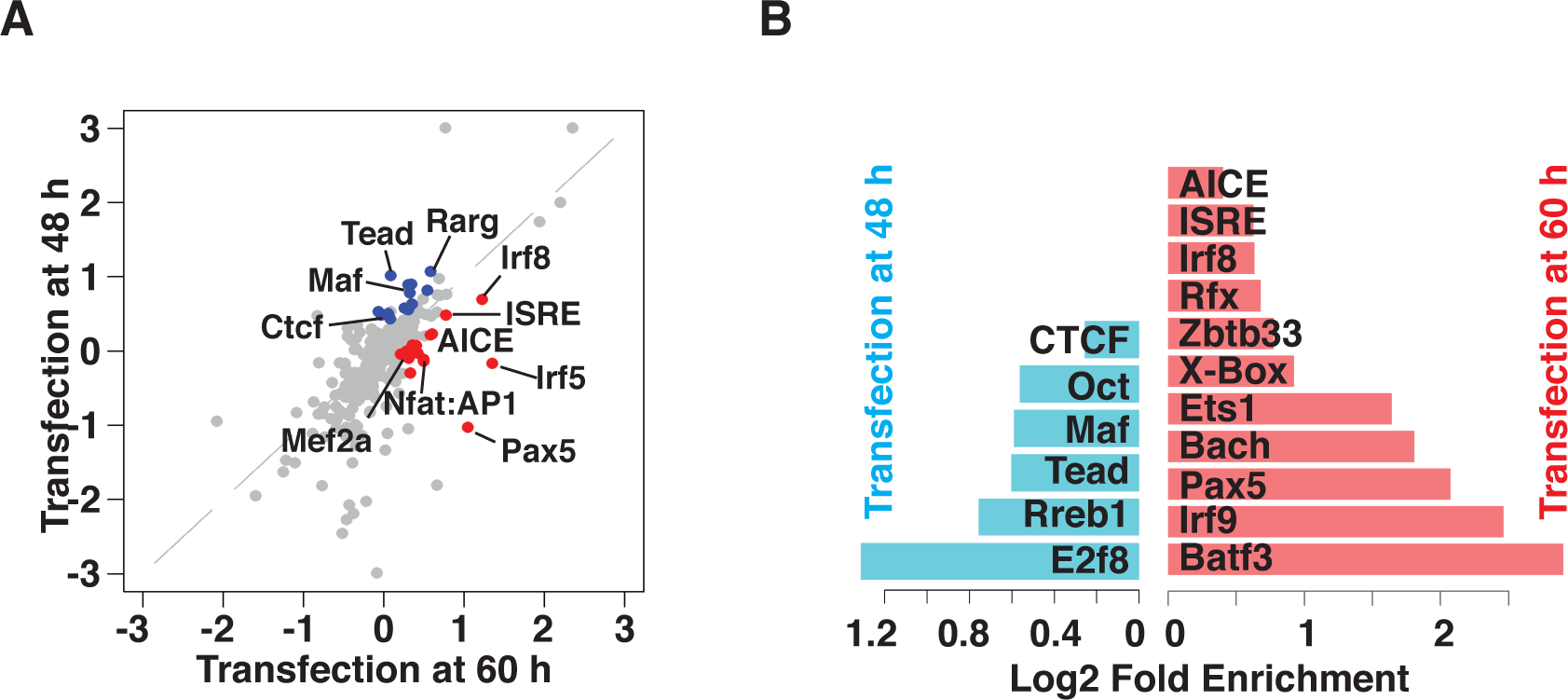
TF motif enrichment analysis of kinetically distinguishable enhancers. **A.** TF motif enrichment analysis (HOMER) was performed using STARR-seq tags for peaks that were uniquely present in the 48h or 60h transfection datasets. TF motif enrichment values (log2 odds ratios) were determined for each transfection time point using the shared peaks for both time points as the background set. In the plotted data, TF motifs that were enriched with logP values < −20 are indicated as colored circles. **B**. TF motif enrichment analysis (HOMER) was performed using STARR-seq tags for peaks that were uniquely present in the 48h transfection dataset using the 60h dataset as background or vice versa. TFs whose motifs (simple or composite elements) are highly enriched (logP < −20) are displayed in the bar plot with their log2 odds ratios.

## References

Ansel, K.M., Lee, D.U., and Rao, A. (2003). An epigenetic view of helper T cell differentiation. Nat Immunol 4, 616–623.

Arnold, C.D., Gerlach, D., Stelzer, C., Boryn, L.M., Rath, M., and Stark, A. (2013). Genome-wide quantitative enhancer activity maps identified by STARR-seq. Science 339, 1074–1077.

Babbitt, C.C., Markstein, M., and Gray, J.M. (2015). Recent advances in functional assays of transcriptional enhancers. Genomics 106, 137–139.

Bhattacharyya, S., Deb, J., Patra, A.K., Thuy Pham, D.A., Chen, W., Vaeth, M., Berberich-Siebelt, F., Klein-Hessling, S., Lamperti, E.D., Reifenberg, K., et al. (2011). NFATc1 affects mouse splenic B cell function by controlling the calcineurin--NFAT signaling network. J Exp Med 208, 823–839.

Braun, T., and Gautel, M. (2011). Transcriptional mechanisms regulating skeletal muscle differentiation, growth and homeostasis. Nat Rev Mol Cell Biol 12, 349–361.

Claussnitzer, M., Dankel, S.N., Kim, K.H., Quon, G., Meuleman, W., Haugen, C., Glunk, V., Sousa, I.S., Beaudry, J.L., Puviindran, V., et al. (2015). FTO Obesity Variant Circuitry and Adipocyte Browning in Humans. N Engl J Med 373, 895–907.

Cobaleda, C., Schebesta, A., Delogu, A., and Busslinger, M. (2007). Pax5: the guardian of B cell identity and function. Nat Immunol 8, 463–470.

Dailey, L. (2015). High throughput technologies for the functional discovery of mammalian enhancers: new approaches for understanding transcriptional regulatory network dynamics. Genomics 106, 151–158.

Davidson, E.H. (2010). Emerging properties of animal gene regulatory networks. Nature 468, 911–920.

Dowen, J.M., Fan, Z.P., Hnisz, D., Ren, G., Abraham, B.J., Zhang, L.N., Weintraub, A.S., Schuijers, J., Lee, T.I., Zhao, K., et al. (2014). Control of cell identity genes occurs in insulated neighborhoods in mammalian chromosomes. Cell 159, 374–387.

Fukaya, T., Lim, B., and Levine, M. (2016). Enhancer Control of Transcriptional Bursting. Cell 166, 358–368.

Gaulton, K.J., Nammo, T., Pasquali, L., Simon, J.M., Giresi, P.G., Fogarty, M.P., Panhuis, T.M., Mieczkowski, P., Secchi, A., Bosco, D., et al. (2010). A map of open chromatin in human pancreatic islets. Nat Genet 42, 255–259.

Glimcher, L.H., and Singh, H. (1999). Transcription factors in lymphocyte development--T and B cells get together. Cell 96, 13–23.

Gonen, N., Futtner, C.R., Wood, S., Garcia-Moreno, S.A., Salamone, I.M., Samson, S.C., Sekido, R., Poulat, F., Maatouk, D.M., and Lovell-Badge, R. (2018). Sex reversal following deletion of a single distal enhancer of Sox9. Science.

Grosschedl, R. (2013). Establishment and maintenance of B cell identity. Cold Spring Harb Symp Quant Biol 78, 23–30.

Heintzman, N.D., Stuart, R.K., Hon, G., Fu, Y., Ching, C.W., Hawkins, R.D., Barrera, L.O., Van Calcar, S., Qu, C., Ching, K.A., et al. (2007). Distinct and predictive chromatin signatures of transcriptional promoters and enhancers in the human genome. Nat Genet 39, 311–318.

Heinz, S., Benner, C., Spann, N., Bertolino, E., Lin, Y.C., Laslo, P., Cheng, J.X., Murre, C., Singh, H., and Glass, C.K. (2010). Simple combinations of lineage-determining transcription factors prime cis-regulatory elements required for macrophage and B cell identities. Mol Cell 38, 576–589.

Hnisz, D., Abraham, B.J., Lee, T.I., Lau, A., Saint-Andre, V., Sigova, A.A., Hoke, H.A., and Young, R.A. (2013). Super-enhancers in the control of cell identity and disease. Cell 155, 934–947.

Ho, J.W., Jung, Y.L., Liu, T., Alver, B.H., Lee, S., Ikegami, K., Sohn, K.A., Minoda, A., Tolstorukov, M.Y., Appert, A., et al. (2014). Comparative analysis of metazoan chromatin organization. Nature 512, 449–452.

Johnson, K.R., Gagnon, L.H., Tian, C., Longo-Guess, C.M., Low, B.E., Wiles, M.V., and Kiernan, A.E. (2018). Deletion of a Long-Range Dlx5 Enhancer Disrupts Inner Ear Development in Mice. Genetics 208, 1165–1179.

Kieffer-Kwon, K.R., Tang, Z., Mathe, E., Qian, J., Sung, M.H., Li, G., Resch, W., Baek, S., Pruett, N., Grontved, L., et al. (2013). Interactome maps of mouse gene regulatory domains reveal basic principles of transcriptional regulation. Cell 155, 1507–1520.

Kim, U., Qin, X.F., Gong, S., Stevens, S., Luo, Y., Nussenzweig, M., and Roeder, R.G. (1996). The B-cell-specific transcription coactivator OCA-B/OBF-1/Bob-1 is essential for normal production of immunoglobulin isotypes. Nature 383, 542–547.

Kurosaki, T., Shinohara, H., and Baba, Y. (2010). B cell signaling and fate decision. Annu Rev Immunol 28, 21–55.

Long, M., de Souza, S.J., and Gilbert, W. (1995). Evolution of the intron-exon structure of eukaryotic genes. Curr Opin Genet Dev 5, 774–778.

Mansson, R., Welinder, E., Ahsberg, J., Lin, Y.C., Benner, C., Glass, C.K., Lucas, J.S., Sigvardsson, M., and Murre, C. (2012). Positive intergenic feedback circuitry, involving EBF1 and FOXO1, orchestrates B-cell fate. Proc Natl Acad Sci U S A 109, 21028–21033.

Martello, G., and Smith, A. (2014). The nature of embryonic stem cells. Annu Rev Cell Dev Biol 30, 647–675.

Mayran, A., and Drouin, J. (2018). Pioneer transcription factors shape the epigenetic landscape. J Biol Chem.

Muerdter, F., Boryn, L.M., Woodfin, A.R., Neumayr, C., Rath, M., Zabidi, M.A., Pagani, M., Haberle, V., Kazmar, T., Catarino, R.R., et al. (2018). Resolving systematic errors in widely used enhancer activity assays in human cells. Nat Methods 15, 141–149.

Nakahashi, H., Kieffer Kwon, K.R., Resch, W., Vian, L., Dose, M., Stavreva, D., Hakim, O., Pruett, N., Nelson, S., Yamane, A., et al. (2013). A genome-wide map of CTCF multivalency redefines the CTCF code. Cell Rep 3, 1678–1689.

Palozola, K.C., Donahue, G., Liu, H., Grant, G.R., Becker, J.S., Cote, A., Yu, H., Raj, A., and Zaret, K.S. (2017). Mitotic transcription and waves of gene reactivation during mitotic exit. Science 358, 119–122.

Rao, S.S., Huntley, M.H., Durand, N.C., Stamenova, E.K., Bochkov, I.D., Robinson, J.T., Sanborn, A.L., Machol, I., Omer, A.D., Lander, E.S., et al.(2014).A 3D map of the human genome at kilobase resolution reveals principles of chromatin looping. Cell 159, 1665– 1680.

Ren, G., Jin, W., Cui, K., Rodrigez, J., Hu, G., Zhang, Z., Larson, D.R., and Zhao, K. (2017). CTCF-Mediated Enhancer-Promoter Interaction Is a Critical Regulator of Cell-to-Cell Variation of Gene Expression. Mol Cell 67, 1049–1058 e1046.

Roy, A.L., Sen, R., and Roeder, R.G. (2011). Enhancer-promoter communication and transcriptional regulation of Igh. Trends Immunol 32, 532–539.

Singh, H., Medina, K.L., and Pongubala, J.M. (2005). Contingent gene regulatory networks and B cell fate specification. Proc Natl Acad Sci U S A 102, 4949–4953.

Swami, M. (2010). Transcription: Shadow enhancers confer robustness. Nat Rev Genet 11, 454.

Tewhey, R., Kotliar, D., Park, D.S., Liu, B., Winnicki, S., Reilly, S.K., Andersen, K.G., Mikkelsen, T.S., Lander, E.S., Schaffner, S.F., et al. (2016). Direct Identification of Hundreds of Expression-Modulating Variants using a Multiplexed Reporter Assay. Cell 165, 1519–1529.

Thompson, D., Regev, A., and Roy, S. (2015). Comparative analysis of gene regulatory networks: from network reconstruction to evolution. Annu Rev Cell Dev Biol 31, 399– 428.

Thurman, R.E., Rynes, E., Humbert, R., Vierstra, J., Maurano, M.T., Haugen, E., Sheffield, N.C., Stergachis, A.B., Wang, H., Vernot, B., et al. (2012). The accessible chromatin landscape of the human genome. Nature 489, 75–82.

Turner, M.L., Hawkins, E.D., and Hodgkin, P.D. (2008). Quantitative regulation of B cell division destiny by signal strength. J Immunol 181, 374–382.

Vaidyanathan, B., Chaudhry, A., Yewdell, W.T., Angeletti, D., Yen, W.F., Wheatley, A.K., Bradfield, C.A., McDermott, A.B., Yewdell, J.W., Rudensky, A.Y., et al. (2017). The aryl hydrocarbon receptor controls cell-fate decisions in B cells. J Exp Med 214, 197–208.

Visel, A., Blow, M.J., Li, Z., Zhang, T., Akiyama, J.A., Holt, A., Plajzer-Frick, I., Shoukry, M., Wright, C., Chen, F., et al. (2009). ChIP-seq accurately predicts tissue-specific activity of enhancers. Nature 457, 854–858.

Vockley, C.M., D’Ippolito, A.M., McDowell, I.C., Majoros, W.H., Safi, A., Song, L., Crawford, G.E., and Reddy, T.E. (2016). Direct GR Binding Sites Potentiate Clusters of TF Binding across the Human Genome. Cell 166, 1269–1281 e1219.

Waardenberg, A.J., Ramialison, M., Bouveret, R., and Harvey, R.P. (2014). Genetic networks governing heart development. Cold Spring Harb Perspect Med 4, a013839.

Weirauch, M.T., Yang, A., Albu, M., Cote, A.G., Montenegro-Montero, A., Drewe, P., Najafabadi, H.S., Lambert, S.A., Mann, I., Cook, K., et al. (2014). Determination and inference of eukaryotic transcription factor sequence specificity. Cell 158, 1431–1443.

Whyte, W.A., Orlando, D.A., Hnisz, D., Abraham, B.J., Lin, C.Y., Kagey, M.H., Rahl, P.B., Lee, T.I., and Young, R.A. (2013). Master transcription factors and mediator establish super-enhancers at key cell identity genes. Cell 153, 307–319.

Xu, H., Chaudhri, V.K., Wu, Z., Biliouris, K., Dienger-Stambaugh, K., Rochman, Y., and Singh, H. (2015). Regulation of bifurcating B cell trajectories by mutual antagonism between transcription factors IRF4 and IRF8. Nat Immunol 16, 1274–1281.

Zabidi, M.A., Arnold, C.D., Schernhuber, K., Pagani, M., Rath, M., Frank, O., and Stark, A. (2015). Enhancer-core-promoter specificity separates developmental and housekeeping gene regulation. Nature 518, 556–559.

Zaret, K.S., and Carroll, J.S. (2011). Pioneer transcription factors: establishing competence for gene expression. Genes Dev 25, 2227–2241.

## References

Dobin, A., Davis, C.A., Schlesinger, F., Drenkow, J., Zaleski, C., Jha, S., Batut, P., Chaisson, M., and Gingeras, T.R. (2013). STAR: ultrafast universal RNA-seq aligner. Bioinformatics 29, 15–21.

Falcon, S., and Gentleman, R. (2007). Using GOstats to test gene lists for GO term association. Bioinformatics 23, 257–258.

Kim, D., Pertea, G., Trapnell, C., Pimentel, H., Kelley, R., and Salzberg, S.L. (2013). TopHat2: accurate alignment of transcriptomes in the presence of insertions, deletions and gene fusions. Genome Biol 14, R36.

Langmead, B., and Salzberg, S.L. (2012). Fast gapped-read alignment with Bowtie 2. Nat Methods 9, 357–359.

Lawrence, M., Huber, W., Pages, H., Aboyoun, P., Carlson, M., Gentleman, R., Morgan, M.T., and Carey, V.J. (2013). Software for computing and annotating genomic ranges. PLoS Comput Biol 9, e1003118.

Phanstiel, D.H., Boyle, A.P., Araya, C.L., and Snyder, M.P. (2014). Sushi.R: flexible, quantitative and integrative genomic visualizations for publication-quality multi-panel figures. Bioinformatics 30, 2808–2810.

Rao, S.S., Huntley, M.H., Durand, N.C., Stamenova, E.K., Bochkov, I.D., Robinson, J.T., Sanborn, A.L., Machol, I., Omer, A.D., Lander, E.S., et al. (2014). A 3D map of the human genome at kilobase resolution reveals principles of chromatin looping. Cell 159, 1665–1680.

Vanhille, L., Griffon, A., Maqbool, M.A., Zacarias-Cabeza, J., Dao, L.T., Fernandez, N., Ballester, B., Andrau, J.C., and Spicuglia, S. (2015). High-throughput and quantitative assessment of enhancer activity in mammals by CapStarr-seq. Nat Commun 6, 6905.

Weirauch, M.T., Yang, A., Albu, M., Cote, A.G., Montenegro-Montero, A., Drewe, P., Najafabadi, H.S., Lambert, S.A., Mann, I., Cook, K., et al. (2014). Determination and inference of eukaryotic transcription factor sequence specificity. Cell 158, 1431– 1443.

